# Agouti and BMP signaling drive a naturally occurring fate conversion of melanophores to leucophores in zebrafish

**DOI:** 10.1101/2024.11.20.624586

**Authors:** Delai Huang, Emaan Kapadia, Yipeng Liang, Leah P. Shriver, Shengkun Dai, Gary J. Patti, Bruno M. Humbel, Vincent Laudet, David M. Parichy

## Abstract

The often-prominent pigment patterns of vertebrates are varied in form and function and depend on several types of pigment cells derived from embryonic neural crest or latent stem cells of neural crest origin. These cells and the patterns they produce have been useful for uncovering features of differentiation and morphogenesis that underlie adult phenotypes, and they offer opportunities to discover how patterns and the cell types themselves have diversified. In zebrafish, a body pattern of stripes arises by self organizing interactions among three types of pigment cells. Yet these fish also exhibit white ornamentation on their fins that depends on the transdifferentiation of black melanophores to white cells, “melanoleucophores.” To identify mechanisms underlying this conversion we used ultrastructural, transcriptomic, mutational and other approaches. We show that melanophore– melanoleucophore transition depends on regional BMP signals transduced through non-canonical receptors (Rgmb-Neo1a-Lrig2) as well as BMP-dependent signaling by Agouti genes, *asip1* and *asip2b*. These signals lead to expression of transcription factor genes including *foxd3* and *runx3* that are necessary to induce loss of melanin by an autophagy-like process, curtail new melanin production, and deploy a pathway for accumulating guanine crystals that, together, confer a white phenotype. These analyses uncover an important role for positional information in specifying ornamentation in zebrafish and show how tissue environmental cues and a novel gene regulatory program have allowed terminal addition of a distinct phenotype to a pre-existing cell type.

**Significance:** Fish often have striking color patterns with important functions in behavior. In zebrafish, the familiar striped pattern forms through self-organizing interactions between pigment cells, yet the white highlights on their fins arise differently—through the transformation of black pigment cells into white ones. This study reveals how this dramatic cell transformation happens: signals from the surrounding tissue, specifically BMP and Agouti proteins, instruct black cells to change their fate. These signals trigger expression of specific genes that cause the cells to break down their black pigment while acquiring white, crystal-like structures. This work shows how local signals in tissues can drive the development of ornamental features and provides insights into how new cell types evolve.

## Introduction

Vertebrate pigmentation has long been a useful system for elucidating mechanisms of pattern formation, particularly in the context of adult traits having behavioral significance and clear selective consequences (1). Given the origins of vertebrate skin pigment cells in embryonic neural crest cells and post-embryonic neural crest derived stem cells—and the many and varied cell types to which these progenitors contribute—studies of pigmentation also can inform our understanding of cell lineage diversification and cell type evolution.

Mammals and birds have a single skin pigment cell, the melanocyte, which can transfer melanin-containing melanosomes to keratinocytes for deposition in hair or feathers (2). Ectothermic vertebrates have multiple types of pigment cells that retain their pigmentary materials intracellularly: black melanophores, with melanin; yellow or orange xanthophores and red erythrophores, with pteridines, carotenoids or both; and iridescent iridophores, with crystalline deposits of guanine in stacks of reflecting platelets (3). In the teleost most studied for its pigmentation, zebrafish *Danio rerio*, interactions among chromatophores generate and maintain a pattern of dark and light horizontal stripes (4), with features of pattern formation resembling a Turing process (5). Orientation of this pattern is thought to depend broadly on the tissue environment—anatomical boundaries and the dynamics of growth (4, 6). Yet the self-organizing properties of pigment cell interactions are revealed by perturbations, with cellular responses that regenerate, reposition or even reorient the pattern (7, 8).

Other pattern features, and pigment cell types, also occur in ectotherms. An example in zebrafish itself is ornamentation of the dorsal fin edge and caudal fin tips, where populations of white pigment cells—leucophores—are situated (Fig. 1*A*,*B*). These highly reflective cells are prominent when fins are flared during behavioral interactions and their presence impacts the propensity of fish to shoal with one another in the laboratory (9). Remarkably, these cells arise from melanophores that lose their melanin over just a few days while acquiring crystalline deposits of guanine, similar chemically to those of iridophores, yet lacking the disc-like structure or spatial organization of reflecting platelets and thereby conferring a white rather than iridescent appearance. In contrast to stripe pattern, white ornaments and the “melanoleucophores” they comprise are unperturbed in mutants that lack xanthophores or iridophores, indicating pattern-forming mechanisms other than self-organizing pigment cell interactions.

**Figure 1.**
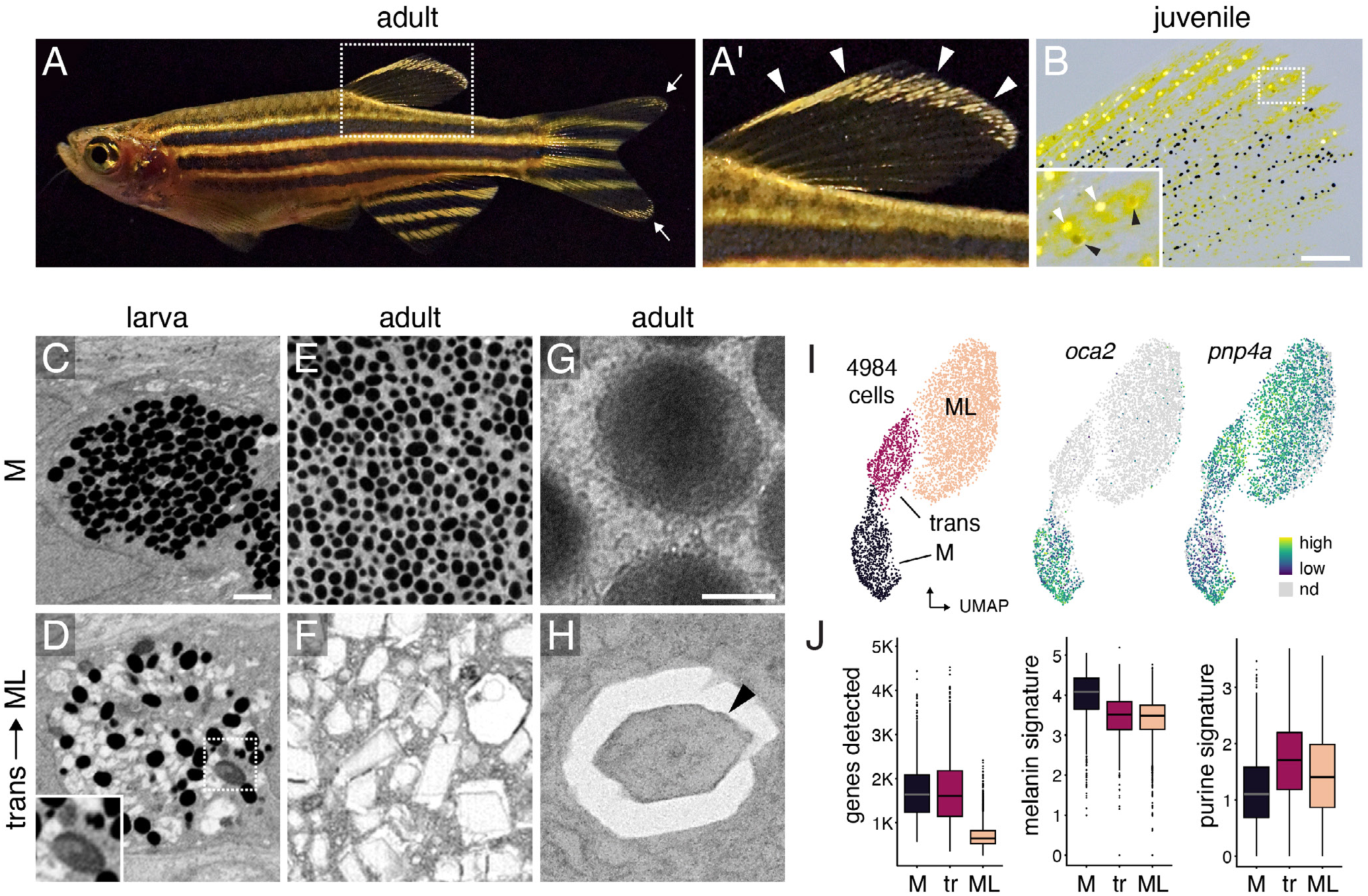
Melanoleucophore (ML) ornamentation and characteristics. (*A*,*A’*) ML occur along anterior and distal edges of the dorsal fin (arrowheads, *A’*) and in lesser numbers at the tips of the caudal fin (arrow, *A*). (*B*) Juvenile fish with white ML (white arrowheads, inset) and grey transitional cells (black arrowheads). Pigmentary organelles have been contracted towards cell centers by epinephrine treatment; yellow pigments are contained within xanthophores (9). (*C,D*) In late larvae, melanophores (M) contain rounded melanosomes whereas transitional cells have heterogeneous pigmentary organelles including irregular or enlarged melanosomes (inset). (*E,F*) In adults, melanophores contain rounded melanosomes whereas ML contain jagged leucosomes with crystalline contents. (*G,H*) Melanosome and leucosome with crystal (arrowhead). *C*–*F*, pigmentary organelles contracted to cell centers by epinephrine treatment, with imaging by focused ion beam scanning electron microscopy (FIB-SEM); *G*,*H*, transmission EM. (*I*) Single cell transcriptomes in UMAP space with cell states (left) and transcript abundances for melanogenesis gene *oca2* and purine salvage gene *pnp4a* (right). (*J*) Compared to melanophores, ML had fewer genes detected, fewer transcripts of genes for melanogenesis, and more transcripts of genes for purine synthesis. Boxplots, medians and interquartile ranges. (Scale bars: *B* 200 µm, 8*C* for *C–F* 1 µm, *G* for *G,H* 0.2 µm).

We sought to better understand the conversion of melanophores to melanoleucophores (ML) in the dorsal fin and whether these events are triggered by localized cues in the tissue environment. Here, we show that melanin loss and guanine accumulation are distinct processes dependent on spatially localized BMP and Agouti signals acting through Foxd3 and Runx3 transcription factors. The result is an overt cellular phenotype and an underlying transcriptomic state markedly different from melanophores, from which ML arise, as well as iridophores. Our analyses demonstrate an important role for positional information in specifying the site and cell state changes that underlie pigmentary ornamentation in zebrafish and show how terminal addition of a new fate can lead to cell type diversification in a vertebrate.

## Results

### Melanosome Degradation, Leucosome Appearance and Independence of Organelle Biogenesis

Melanin and melanosomes are stable under a variety of conditions and persist when melanogenesis is inhibited (10, 11). We therefore hypothesized that rapid (∼4 d) loss of melanin during the transition of melanophores to melanoleucophores (ML)(9) results from degradative processes. Consistent with organellar breakdown (12), transitional cells contained melanosomes of irregular shapes, sizes, and densities, often associated with endosomal bodies (Fig. 1*D*; *SI Appendix* Fig. S1A–C), and contrasting with the more homogeneous melanosomes of melanophores (Fig. 1*C,E,G*; *SI* Movies S1–S3). We did not detect melanosomes in extracellular spaces (*SI* Movie S2), suggesting loss of melanin during the melanophore–ML transition is not accomplished by shedding as might be hypothesized by comparison with mammalian melanocytes (13). Transitional cells also contained small and rounded presumptive “leucosomes” of various shapes and sizes (Fig. 1*D*; *SI Appendix* Fig. S1A–C). ML of adult fish contained larger, jagged leucosomes bearing crystalline material (Fig. 1*F,H*) but also small numbers of irregular melanosomes similar to those of transitional cells (*SI Appendix* Fig. S1D, *SI* Movie S4).

ML contain crystals of β-guanine (9) (*SI Appendix* Fig. S1E; *SI* Table S1). Since iridophores also contain crystals of β-guanine (14, 15), we hypothesized that iridophores and ML share genetic requirements for guanine accumulation. To test this idea, we focused on *pnp4a*, which encodes a purine nucleoside phosphorylase (PNP) of the purine salvage pathway expressed in both cell types (9). When targeted by CRISPR/Cas9 to generate premature termination codon (PTC) alleles, we found that homozygotes had reduced iridescence of iridophores and lower leucosome/guanine contents of ML, indicating overlapping requirements between cell types (*SI Appendix* Figs. S2,3*A,C*). Electron microscopy and the phenotype of *pnp4a* mutants further suggested that melanosome and leucosome biogenesis are likely to be distinct from one another. To confirm this inference, we assessed leucosome contents in mutants for *oca2*, encoding a channel that regulates melanosomal pH and activity of the melanogenesis-rate limiting enzyme Tyrosinase (16, 17), as well as *pmela*, encoding a protein essential for melanosomal matrix deposition (18, 19). Despite reduced or absent melanin, *oca2* and *pmela* PTC alleles had normal leucosome/guanine contents (*SI Appendix* Figs. S2,3*A–C*), consistent with distinct processes of organellogenesis.

### Transcriptomic Signatures of Cell Type Conversion

We further assessed cell state change by single cell RNA-Sequencing (scRNA-Seq) after isolating *tyrp1b*:PALM-mCherry+ (9) cells of the melanophore–ML lineage from juvenile fish. Unsupervised clustering revealed cell groups identifiable as melanophores, transitional, or ML, with trajectory analyses bridging states in pseudotime and representing an inferred pathway of differentiation concordant with previous fate mapping *in vivo* (Fig. 1*I*; *SI Appendix* Figs. S4*A*–*F*; *SI* Table S2–4) (9). These analyses revealed that ML express many fewer genes than melanophores or transitional cells (Fig. 1*J*), with cells at later steps in pseudotime having fewer genes detected than cells at earlier steps, even when sequenced to similar depth (*SI Appendix* Figs. S4*F*,*G*). ML likewise had reduced signatures of transcriptional and translational activity and cell cycle genes (SI Appendix Fig. S4H). Transitional cells did not upregulate neural crest marker expression overall, suggesting that melanophore–ML transition does not entail dedifferentiation towards an unspecified progenitor state (*SI Appendix* Fig. S4H; *SI* Table S6).

As anticipated, transitional cells and ML expressed melanogenesis genes at lower levels than melanophores (Fig. 1*I*,*J*) yet profiles varied markedly among genes: only *oca2* had a distinct boundary in expression, suggesting its downregulation may be essential for curtailing new melanin production (*SI Appendix* Fig. S4I). Transcripts of purine pathway genes were more abundant in transitional cells (Fig. 1*N*; *SI Appendix* Fig. S4I). These cells continued to express *melanocyte inducing transcription factor a* (*mitfa*), the master regulator of melanophore development, and *kita*, encoding a receptor tyrosine kinase essential for melanophore differentiation and morphogenesis (2), both having essential functions in ML (9) (*SI Appendix* Fig. S4I). Pathway enrichment analyses confirmed a strong signature of MAPK signaling in ML, consistent with Kita activity, and suggested roles for an autophagy-like process in cellular remodeling (*SI Appendix* Fig. S5A,*B*, Table S7). Given this observation and indications of melanosome degradation, we predicted that blockade of autophagy mechanisms should lead to persisting melanin, and indeed fish treated with the autophagosome-lysosome fusion inhibitor bafilomycin A1 retained melanin in otherwise white cells (*SI Appendix* Fig. S5C).

The seemingly large-scale differences in transcriptomic state and cytoarchitecture between melanophores and ML led us to ask how these alterations compare to differences within and between other pigment cell lineages. To address this question we used similarly obtained scRNA-Seq data for pigment cells on the body of post-embryonic zebrafish (20). In these comparisons, M–ML differences were not less than those between undifferentiated body progenitors and melanophores, nor between body melanophores and xanthophores, despite ML arising directly from melanophores (*SI Appendix* Fig. S6A*,B*). Though ML differentiation entailed purine synthesis gene upregulation (Fig. 1*J*), similar to iridophores, neither transitional cells nor ML expressed genes for Tfec, a transcription factor essential for iridophore specification (21) or Ltk, a receptor tyrosine kinase essential for iridophore differentiation and morphogenesis (22) (*SI Appendix* Fig. S6C). Thus, ML depend on a different gene regulatory program than iridophores, despite their pigmentary phenotype relying on a common structural feature— guanine crystals.

### *asip1* and *asip2b* Together Induce Melanophore to ML Transition

We sought to identify factors that instruct or permit the melanophore–ML transition. Given the limited anatomical distribution of ML, we hypothesized the presence of positional cues anterodorsally in the fin, and so predicted that genes encoding one or more secreted factors should be expressed at higher levels along this fin edge as compared to basally. By scRNA-Seq of whole fin, 592 genes were expressed at higher levels (*q*<0.05) at the fin edge, amongst which *agouti signaling protein 1* (*asip1*) had the largest difference in transcript abundance (Fig. 2*A*; *SI Appendix* Fig. S7A*,B*; *SI* Tables S8,9). Asip1 and the orthologous protein of mammals, Asip, are Melanocortin-1 (Mc1r) receptor antagonists and their ventral expression inhibits differentiation of melanophores and melanocytes, respectively, resulting in light bellies (23–26). In fin, *asip1* was expressed by extracellular matrix producing, *basonuclin-2*+ dermal mesenchymal cells (Fig. 2*B,C*; *SI Appendix* Fig. S7C*,D*; *SI* Table S9), similar to hypodermal cells that support chromatophores on the body (27). Melanocortin receptor genes were, correspondingly, expressed by melanophores–ML (*SI Appendix* Fig. S7F). To test roles for Asip1 signaling we generated *asip1* and *mc1r* mutants. *asip1* mutants had mild perturbations to fin melanophore pattern, but ML complements equivalent to wild-type (Fig. 2*D*,*E*). By contrast, *mc1r* mutants had extra ML that covered a greater proportion of the fin compared to wild-type, consistent with a requirement for Mc1r repression in ML development (*SI Appendix* Figs. S2, S7*G*; *SI* Table S10).

**Figure 2.**
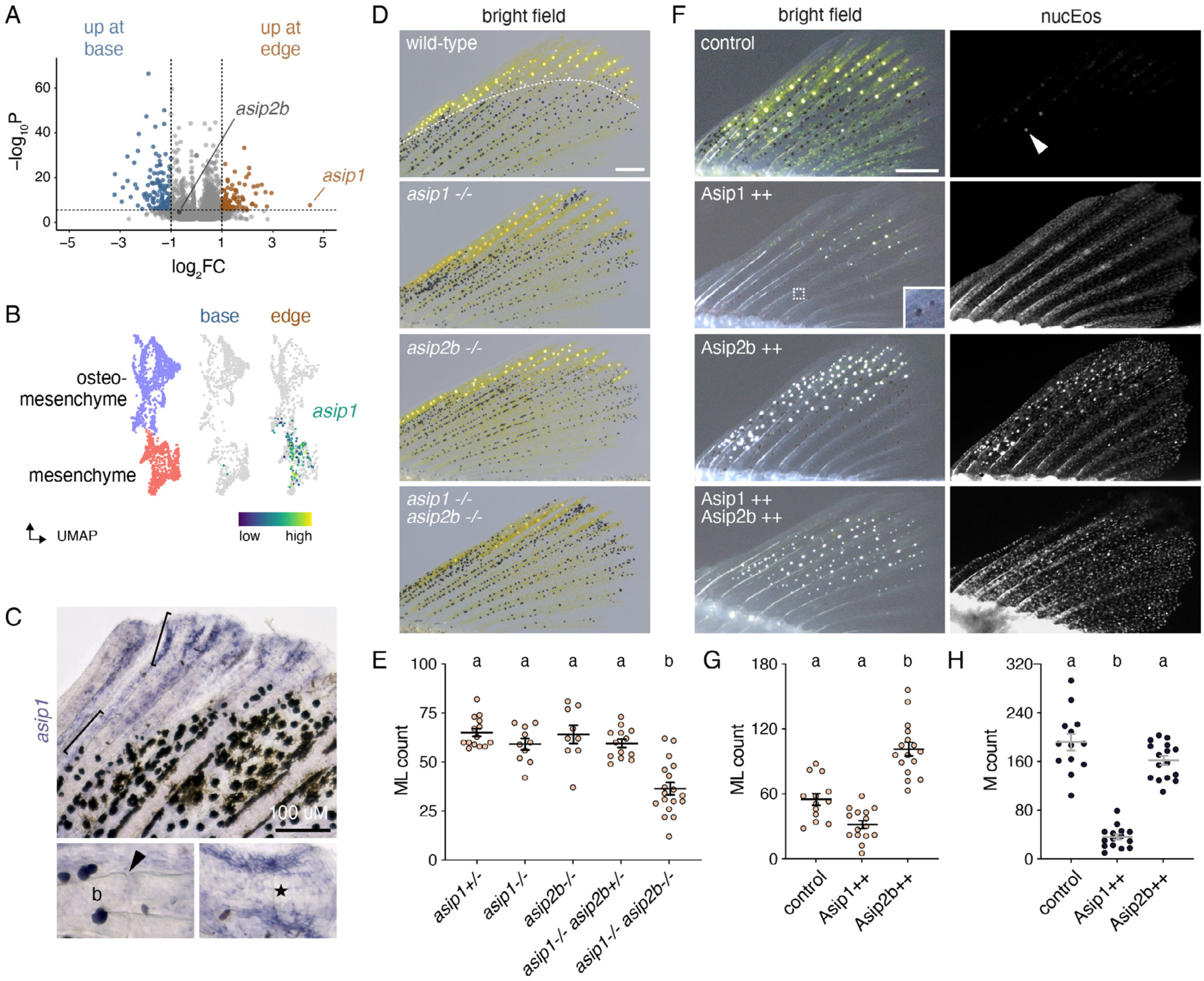
Agouti genes contribute to ML development. (*A*) Among cells in the melanophore–ML environment (dermis, osteoblasts, osteomesenchyme, epidermis, neurons, glia; *n*=3172 cells) *asip1* was expressed 4.5-fold higher in the fin edge than base (*p*=2.4E-8, *q*=0.0004). Transcripts for a second Agouti gene, *asip2b*, were present, but not differentially abundant. The plane along which fins were dissected into edge vs. base for scRNA-Seq is indicated by the dashed line in *D* (top). (*B,C*) *asip1* transcripts were limited to mesenchyme and osteomesenchyme in edge (UMAP and cell type colors subsets of *SI Appendix* Fig. S7A). Brackets in *C* correspond to details. Arrowhead, joint between proximal fin bones (b; lepidotrichia). *asip1* transcripts were most abundant in cells adjacent to distal developing lepidotrichia (star). (*D,E*) Fish mutant for *asip1* or *asip2b* developed normal numbers of ML type. By contrast, fish doubly mutant for both loci had significantly fewer ML than wild-type siblings. Overall ANOVA, *F*_4,56_=18.9, *p*<0.0001. Shared letters above groups indicate means not significantly different (*p*>0.05) by *post hoc* Tukey-Kramer HSD. (*F–H*) Asip1 overexpression led to fewer ML, and melanophores, whereas Asip2b overexpression resulted in more ML, without significantly affecting melanophore numbers. Inset for Asip1++ shows two lightly melanized melanophores. In qualitative results from fish that expressed both Asip1 and Asip2b, ML often developed widely across fins nearly devoid of melanophores. nucEos, widespread expression of nuclear localizing Eos, linked by 2A sequence to Asip1 or Asip2b in *hsp70l*-driven transgenes. Arrowhead, reflecting ML guanine crystals in non-transgenic control. Overall ANOVAs, *G,H*: *F*_2,41_=46.2, 84.6, *p*<0.0001. Bars in *E,G,H*, means±SE. (Scale bars: *C* 100 µm, *D,F* 200 µm).

Due to the teleost specific genome duplication, teleosts have several Agouti-related genes (28–30) so we considered that *asip1* could function with another gene-family member. Indeed, we detected transcripts of a second Mc1r antagonizing factor, Asip2b (previously, Agouti related peptide 2, Agrp2)(31) in mesenchyme and osteomesenchyme along the fin base and also at the fin edge and so generated a PTC allele for this locus as well (Fig. 2*A*; *SI Appendix* Figs. S2, S7*C–E*). Fish mutant for *asip2b* had normal ML complements, whereas fish doubly mutant for *asip1* and *asip2b* developed significantly fewer ML (Fig. 2*D,E*). Given this finding we asked whether Agouti genes were also sufficient to drive ML numbers in excess of the wild type (Fig. 2*F–H*). Transgenic fish expressing heat-shock (*hsp70l*) inducible Asip1 developed fewer ML, but also fewer melanophores, suggesting ML deficiency resulted from loss of melanophore progenitors. By contrast, fish transgenic for heat-shock inducible Asip2b developed excess ML, with a corresponding, albeit non-significant reduction in melanophores. Fish mosaically expressing both transgenes developed extra ML, yet very few melanophores. Together, these results supported a model in which ML development requires synergistic activities of *asip1* and *asip2b* where their expression overlaps at the fin edge.

### Bmp Signaling Promotes Melanophore–ML Transition

Even fish doubly mutant for *asip1* and *asip2b* retained some ML indicating essential roles for additional factors as well. Several pathways contribute to fin patterning and morphogenesis and would be seemingly well-positioned for such a role. To test this idea, we examined Wnt and Bmp pathways, which function in other contexts in both fin (32–34) and pigmentation (35–38). Several Wnt ligand genes were expressed at higher levels at the fin edge than base and melanophores–ML expressed Wnt receptor genes (*SI Appendix* Fig. S8A–C; *SI* Table S11). Nevertheless, a Wnt signaling activity reporter was not detectably expressed by these cells (*SI Appendix* Fig. S8D) and inhibition of Wnt signaling did not affect ML number independently of fin outgrowth (*SI Appendix* Fig. S7E; *SI* Table S11).

Consistent with differences in BMP signaling across the fin, several genes encoding Bmp ligands or inhibitors were differentially expressed between base and edge (*SI Appendix* Fig. S9A; *SI* Table S11). Suggesting competence to receive BMP signals, melanophores–ML expressed several BMP/TGFβ receptor genes (*SI Appendix* Fig. S9B; *SI* Table S11). Indeed, BMP signaling activity assayed by anti-pSmad immunoreactivity was apparent in melanophores, transitional cells and ML, and its strength was correlated with proximity to the fin edge (Fig. 3*A–C*). To test roles for Bmp signaling we treated larvae with BMP pathway inhibitor LDN-193189 at a dose permissive for fin outgrowth, which resulted in fewer ML and more melanophores than vehicle (DMSO) control larvae (Fig. 3*D,E*; *SI* Table S12). Other *Danio* species also have ML (9)(*SI Appendix* Fig. S10) and an evolutionarily conserved function for BMP signaling in ML development was indicated by a corresponding effect of BMP inhibition in *D. albolineatus* (Fig. 3*D,E*).

**Figure 3.**
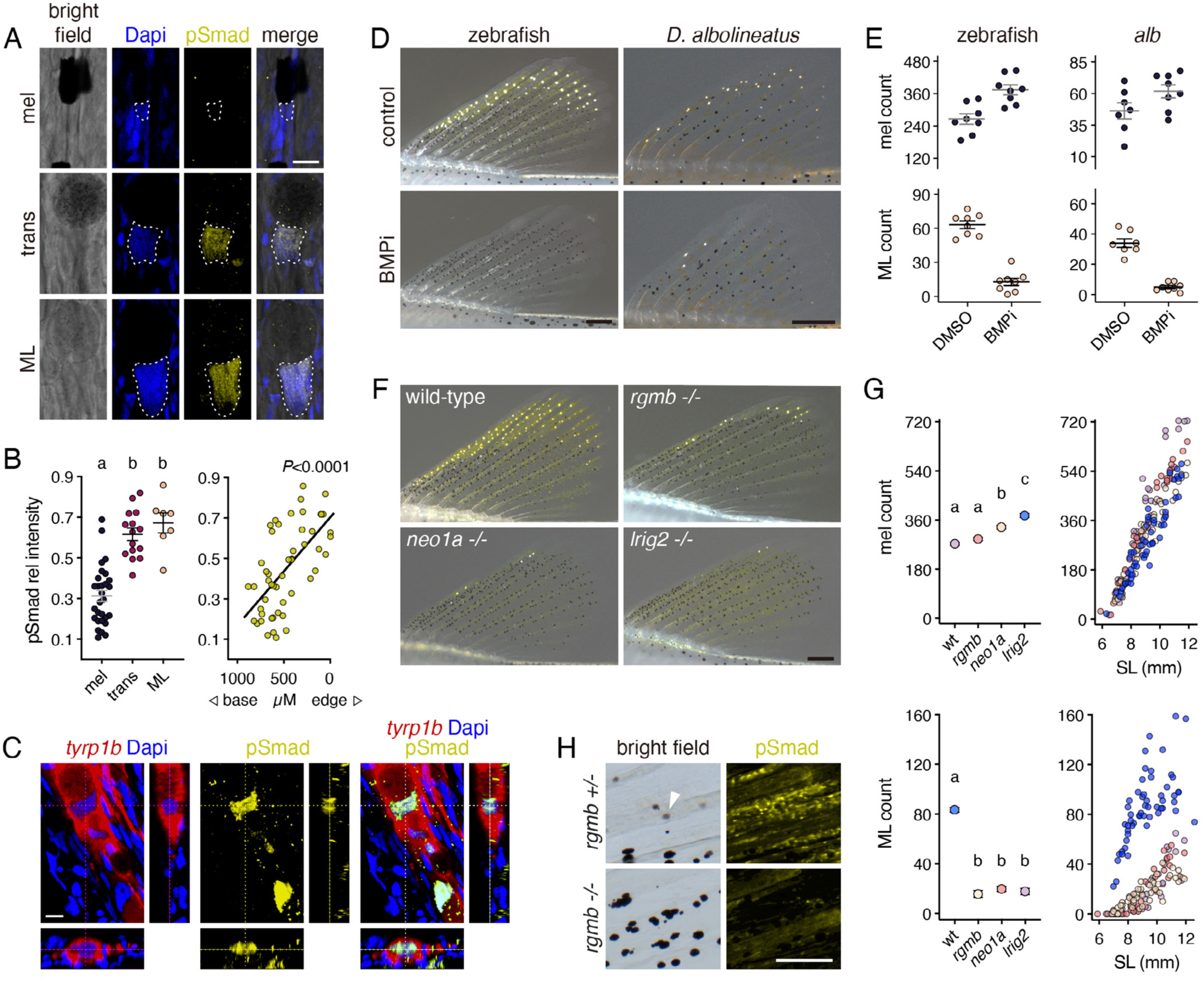
BMP-dependent melanophore–ML transition involves *rgmb-neo1a-lrig2*. (*A*) pSmad immunoreactivity was prominent in nuclei (outlined) of transitional cells and ML. (*B*) Intensities of pSmad staining relative to Dapi across cell states (left) and locations (measured as distance from the most distal ML; *R*^2^=0.39). Shared letters indicate means not significantly different in Tukey-Kramer *post hoc* comparisons (overall *F*_2,48_=35.15, *P*<0.0001). (*C*) ML expressing membrane-targeted *tyrp1b*:PALM-mCherry, illustrating pSmad immunoreactivity in Dapi-labeled nucleus. (*D,E*) BMP signaling inhibitor (BMPi) led to zebrafish and *D. albolineatus* having marginally greater numbers of melanophores (*F*_1,14_=16.59, *P*<0.005 and *F*_1,13_=3.58, *P*=0.08, respectively) and fewer ML (*F*_1,14_=98.59, *P*<0.0001 and *F*_1,13_=113.83, *P*<0.0001). (*F*) Mutants for premature termination alleles of *rgmb*, *neo1a* and *lrig2* had more melanophores and were severely deficient for ML. (*G*) Melanophores (upper) and ML (lower) across genotypes. Left plots show least square means (±SE) after controlling for variation in standard length (SL), a proxy for stage of development (66). Shared letters indicate means not significantly different in *post hoc* comparisons after controlling for standard length and genotype x standard length interactions (all factors in ANOVAs, *P*<0.0001; numbers of ML square-root transformed to control for increased variance in residuals with larger counts). Fin lengths for a given standard length did not differ among genotypes (*SI Appendix* Fig. S8G). (H) *rgmb* mutants exhibited markedly reduced anti-pSmad immunoreactivity across the fin. Arrowhead, transitional cell (in transmitted illumination). (Scale bars: *D* 200 µM for zebrafish and *D. albolineatus*, *F* 200 µm, *H* 100 µm).

To identify loci required for BMP effects we targeted BMP ligand gene *gdf6a*, expressed by fin mesenchyme and having the largest difference in transcript abundance of ligand genes between fin edge and base (*SI Appendix* Fig. S9A; *SI* Table S7). Mutants homozygous for a PTC allele were semi-viable but individuals that survived had fewer ML and more melanophores than wild-type, even after controlling for reduced fin size (*SI Appendix* Figs. S2, S9*D*; *SI* Table S12).

### Rgmb-Neo1a-Lrig2 Requirement for ML Development

To further evaluate BMP pathway roles, we considered targeting Bmp receptors expressed by melanophores–ML (*SI Appendix* Fig. S9B) but published images of homozygous *bmpr2a* and *bmpr2b* mutants (39) suggested that ML persist in these fish, perhaps owing to functional redundancies. Given expression of canonical BMP receptors across a range of cell-types and the possibility that assessment of ML development could be precluded by pleiotropic defects in fish doubly or triply mutant for such receptors, we considered alternative targets. Among genes known to function in BMP signaling we found that *rgmb*, encoding Repulsive Guidance Molecule b (RGMb) (40, 41), was expressed relatively prominently in melanophores and transitional cells (*SI Appendix* Fig. S9B*,C,E*). Homozygous mutants for PTC alleles of *rgmb* did not have defects in fin growth (*SI Appendix* Figs. S2, S9*F*) but did develop fewer ML than wild-type siblings (Fig. 3*F,G*; *SI* Table S12). RGM proteins have high affinity for BMPs and can enhance BMP signaling in ways still incompletely understood but involving formation and internalization of a cell surface complex in which RGMb forms a bridge between BMP ligands and Neogenin (Neo) (42, 43). Consistent with an RGMb-enhancement of BMP signaling, *rgmb* mutants had reduced anti-pSmad immunoreactivity throughout the fin (Fig. 3*H*). Similarly, *neo1a*, encoding Neogenin 1a, was expressed in melanophores–ML and mutants homozygous for PTC alleles had fewer ML, and also more melanophores, than wild-type (Fig. 3*F,G*; *SI Appendix* Figs. S2, S9*B*,*C*,*E*,*F*, *SI* Table S12). Finally, because Lrig (Leucine-rich repeats and immunoglobulin-like domain) proteins have been implicated in BMP signaling and in protecting Neogenin from proteolytic ectodomain cleavage (44, 45), we asked whether Lrig2 contributes to melanophore–ML transition. *lrig2* was expressed in melanophores and transitional cells and PTC mutants had ML deficits indistinguishable from *rgmb* or *neo1a* mutants, along with an excess of melanophores (Fig. 3*F,G*; *SI Appendix* Figs. S2, S9*B*,*C*,*E*,*F*, *SI* Table S12). These observations support roles for Rgmb acting with Neo1a and Lrig2 to promote BMP signaling and melanophore–ML transition.

### BMP-Dependence of Agouti Signaling

Given roles for BMP and Agouti genes in ML development we asked whether these pathways are interdependent. To assess whether BMP signaling requires Agouti, we examined pSmad staining: immunoreactivity was indistinguishable between wild-type and *asip1* or *asip2b* mutants, as well as between wild-type and fish overexpressing Asip1 and Asip2b (*SI Appendix* Fig. S11A). To determine if Agouti signaling requires BMP, we compared gene expression between wild-type and *rgmb* mutants by *in situ* hybridization. This revealed less staining and more limited distribution of staining for *asip1* and *asip2b* in the BMP signaling deficient *rgmb*−/− background (*SI Appendix* Fig. S11B). Reasoning that reduced Agouti gene expression could reflect a function for BMP in dermal development, we stained for dermal markers *bnc2* and *fmoda*, which were also reduced in *rgmb*−/− (*SI Appendix* Fig. S11B). These observations suggested that BMP signals promote the development of dermal cells and their expression of Agouti genes. To assess functional consequences of such interactions, we asked if effects of Agouti overexpression differed between wild-type and *rgmb* mutants. Asip1 and Asip2b reduced and increased ML numbers, respectively, yet the increase driven by Asip2b was attenuated in the *rgmb* mutant background (*SI Appendix* Fig. S11C, *SI* Table S10). Together these observations imply that BMP promotes expression of Asip1 and Asip2b and potentiates signaling by the latter.

### BMP-Signaling Drives Known and Novel Pigmentation Genes to Effect ML Development

To identify additional mechanisms downstream of BMP signals, we treated wild-type larvae with BMP inhibitor LDN-193189 for 48 h during the period when melanophores transition to ML. We then isolated melanophores and ML (*tyrp1b*:PALM-mCherry+) and cells of the fin environment (mCherry–) for scRNA-Seq (*SI Appendix* Fig. S12A,*B*, Tables S13, S14). Numbers of melanophores and ML did not diverge between treated and control fish during the brief duration of the experiment (*SI Appendix* Fig. S12C, *SI* Table S12). Among mCherry– cells, *asip2b* was expressed more robustly in fin mesenchyme of vehicle control (BMP+) larvae than larvae treated with inhibitor (BMP–; log_2_FC=1.1, *q*=4.9E-09; *SI Appendix* Fig. S12D), consistent with reduced *asip2b* expression in *rgmb* mutants. Among mCherry+ cells, several pigmentary genes were expressed at higher levels in control (BMP+) cells, including *pnp4a* and other genes of purine salvage, *tbx3b*, an orthologue of mammalian *Tbx3*, which has roles in phaeomelanogenesis (46), and *foxd3*, encoding a winged helix transcription factor required to maintain pluripotency of neural crest and in specification of some neural crest derivatives (21, 47–49) (Fig. 4*A*; *SI Appendix* Fig. S12D,*E*, *SI* Table S14). Given the varied roles of *foxd3*, and peak expression in transitional cells (*SI Appendix* Fig. S12F), we sought to determine its functional significance. Homozygous mutants are early lethal so we examined fish mosaic for CRISPR/Cas9 induced somatic mutations. We also tested consequences of transgenic Foxd3 induction. These analyses confirmed a requirement and sufficiency: increasing efficacies of *foxd3* mutagenesis led to fewer ML, whereas Foxd3 overexpression led to more ML (Fig. 4*B*; *SI Appendix* Fig. S12G, *SI* Table S15).

**Figure 4.**
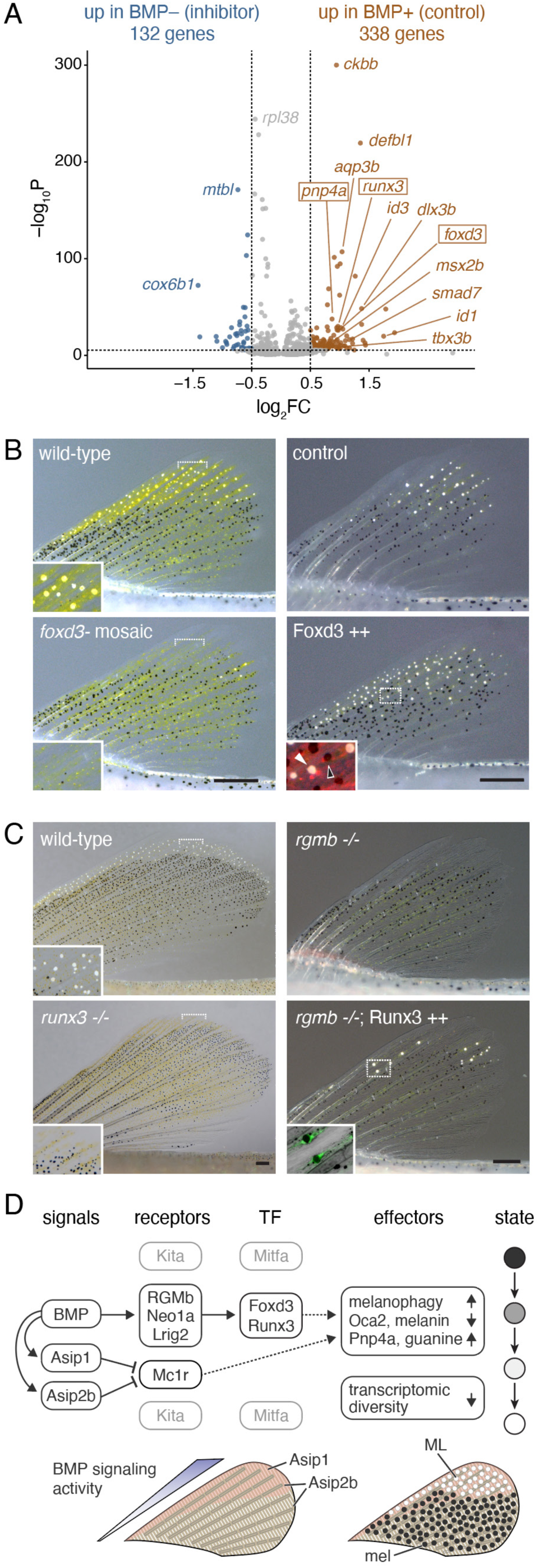
BMP-dependent genes in ML development. (*A*) Differential gene expression in melanophore–ML isolated from acutely inhibitor treated (BMP-) or DMSO-control (BMP+) early juvenile fish. Dotted lines, log_2_FC>|0.5|, *q*<0.05. (*B*) Fish mosaic for somatic mutations in *foxd3* had fewer ML, whereas heat-shock induced Foxd3-2a-nuc-mCherry led to significantly more ML. Insets, details (left) and fluorescence overlay (right) with nuclear mCherry in binucleated Foxd3++ ML (white arrowhead) and melanophore (black arrowhead). (*C*) *runx3* mutants were viable, had normal fins, and lacked nearly all ML (means±SE ML, wild-type: 144±7; *runx3*: 5±8 ML; *F*_1,14_=163.6, *P*<0.0001). Insets, details (left) and overlay showing cytosolic EGFP in ML, with guanine deposits black in this pseudo-brightfield image. (*D*) Upper, Model for major contributors to melanophore– ML conversion. Dark, addressed in this study. Grey, addressed in (10). “BMP” refers to pathway signaling activity overall, as inferred by pSmad immunostaining (*SI Appendix* Fig. S3B), likely to involve ligand Gdf6a (*SI Appendix* Fig. S9D) and possibly other differentially distributed factors (*SI Appendix* Fig. S9A). Dashed lines indicate potential regulatory linkages. Lower, Inferred regions exposed to higher levels of signaling factors during fin outgrowth. ML differentiate where BMP signaling and Agouti gene expression are strongest. Additional key figures: BMP➝Asip1 (*SI Appendix* Fig. S11B); BMP➝Asip2b (*SI Appendix* Figs. S11*B*, S12*D*); BMP➝RGMb/Neo1a/Lrig2 (Fig. 3D–*H*); Asip1, Asip2b gene expression (Figs. 2*A*,*C*, *SI Appendix* S7*C*,*E*, S11*B*); BMP➝➝Foxd3 & Runx3➝➝transitional (Fig. 4A–C, *SI Appendix* Fig. S12G); autophagy-like “melanophagy” (Fig. 1D, *SI Appendix* Figs. S1B, S5, SI); Oca2, melanin & Pnp4a, guanine (Fig. 1I,*J*, *SI Appendix* Figs. S1*E*, S3*A*,*C*); transcriptomic diversity (Fig. 1J, *SI Appendix* Fig. S4G,*F*). (Scale bars: *B*,*C* 200 µm).

Beyond known pigmentary genes, BMP+ cells expressed higher levels of genes encoding factors that function downstream of, or in concert with, BMP in other contexts (e.g., Id1–3, Dlx3, Msx2, Smad7, Runx3). To determine the significance of such loci for ML development we screened 39 additional candidates by CRISPR/Cas9 mutagenesis and found that PTC alleles of one, transcription factor gene *runx3* (*SI Appendix* Fig. S12D,*F*, *SI* Table S16), led to severe ML deficiencies when homozygous, reminiscent of BMP pathway mutants (Fig. 4*C*). We therefore asked if Runx3 was sufficient to rescue ML in *rgmb* mutants; indeed, mosaic Runx3 restored ML when expressed by melanophores–ML (Fig. 4*C*; *SI Appendix* Fig. S12H, *SI* Table S15). Thus, BMP functions through both Foxd3 and Runx3 in promoting ML development.

## Discussion

Our results provide new insights into the development of pigmentary ornamentation in *Danio* fishes. In contrast to other pigment cells of zebrafish and its relatives, ML are limited to the edge of the dorsal fin and the distal tips of the caudal fin. Analyses presented here indicate a role for positional information in the development of ML at the fin edge, and suggest a working model for the differentiation of these cells from melanophores (Fig. 4*D*): (i) WNT and BMP signals promote fin outgrowth and the latter contributes to a landscape of positional information available to pigment cells; (ii) BMP signaling promotes dermal development and expression of Agouti genes *asip1* and *asip2b* by subsets of these cells; (iii) melanophores responsive to BMP signals through Rgmb-Neo1a-Lrig2 and exposed simultaneously to Asip1 and Asip2b initiate a transition to the ML phenotype, upregulating Foxd3 and Runx3 transcription factors; (iv) melanin and melanosomes are degraded by an autophagous process while new melanogenesis is curtailed by downregulation of Oca2 and other factors; (v) a pathway of leucosome biogenesis and purine accumulation is upregulated, depending in part on Pnp4a; and (vi) as a terminal phenotype is acquired, ML reduce gene expression overall to levels markedly less than those of melanophores.

The development of ML from melanophores provides an interesting contrast for genes and pathways that have pigmentary functions in other contexts. For instance, the role of Agouti genes *asip1* and *asip2b* in promoting melanophore–ML transition somewhat resembles roles for these genes separately in repressing melanization ventrally (rodent Asip, zebrafish Asip1) or dorsally (rodent Asip, avian Agouti), or repressing stripe melanophore differentiation laterally (cichlids, Asip2b)(23–25, 28, 50, 51). Here, *asip1* and *asip2b* expression depended at least in part on BMP signaling, concordant with BMP dependence of Asip expression in mouse (52). Coincident deployment of *asip1* and *asip2b* to supply positional information raises interesting questions about gene regulatory mechanisms and their evolution. The different effects of these factors when overexpressed also raise the possibility of non-overlapping downstream mechanisms of signaling independent of Mc1r antagonism, perhaps through a different melanocortin receptor (31).

Roles for BMP signals though Rgmb-Neo-Lrig2 (40, 41) likewise add a new dimension to pigmentary functions of this pathway. BMP signaling in zebrafish biases the fates of progenitor cells away from melanophores and towards iridophores, at least in part by repressing *mitfa* and other genes required for melanogenesis (36). Repressive effects on melanogenesis have been found in mammalian systems as well (52, 53). Yet BMP also promotes postnatal differentiation of WNT-activated hair follicle stem cells as melanocytes, with progenitors accumulating in excess if BMP signaling is blocked (35). Here, pharmacological inhibition and mutant phenotypes indicated an essential role for BMP signals in melanophore–ML transition, with BMP inhibition leading to small but proportionate increases in progenitors—in this case melanophores. The BMP-dependence of Agouti gene expression implies that BMP signals in this system act indirectly on melanophores and ML, and other kinds of indirect effects might be envisaged [*c.f.*,(54)]. Yet direct effects on melanophores–ML also can be inferred, given expression of Rgmb-Neo-Lrig2 and other receptor genes by these cells, the presence of signaling activity revealed by pSmad immunoreactivity specific to these cells, and the correspondence of such activity with anatomical position and fate within this lineage. Whatever the relative contributions of direct and indirect signals, they seem not to involve repression of Mitfa, which is expressed and required through the melanophore–ML transition (9). Comparisons across systems thereby suggest that consequences of BMP signaling for cell fate transitions depend on cellular context including collaborating pathways (e.g., WNT vs. Agouti).

Downstream of BMP, Foxd3 and Runx3 transcription factor genes were upregulated specifically in transitional cells and were required for melanophore–ML transition. Intriguingly both gene products can function as pioneer factors and both can activate or repress transcription depending on context (47, 55). Foxd3 maintains pluripotency of neural crest cells and later contributes to specifying neuronal, glial and iridophore fates over melanocyte or melanophore fates (48, 49, 56) and is regulated by BMP signals in other contexts (57, 58). Runx3 heterodimerizes with Core binding factor subunit β (CBFβ) and interacts with Smads and other factors to promote specific neuronal, immune and skeletal fates. Runx3 has roles in cancers including melanoma but has not previously been ascribed roles in normal pigment cells (59–62). Direct targets of Foxd3 and Runx3—as well as Mitfa—in repressing a melanized phenotype or promoting a leucistic phenotype will be interesting to identify, particularly as they appear to recruit a pathway for guanine crystal biogenesis through mechanisms independent of those used by other guanine crystal containing cells, iridophores (15, 22). In the absence of iridogenic transcription factor Tfec, expression of purine synthesis gene *pnp4a* can be driven by Mitfa, and one might anticipate such a dependency in this context as well (21, 63).

Finally, mechanisms uncovered here shed new light on diversification of cell types and modes of pattern formation (3, 64). We speculate that evolutionary cooptions of *foxd3* and *runx3* expression to the melanophore–ML lineage have been important in the terminal addition of a new fate for a subset of melanophores. Expression of these genes could reflect recruitment of BMP-responsive Agouti gene expression to the fin edge if there were preexisting regulatory linkages between *foxd3* and *runx3* and these signaling pathways, or such linkages may have been acquired in the form of *cis*-regulatory elements conferring responsiveness to BMP, Agouti or both signals, allowing cells to exploit a preexisting landscape of positional information. Whatever the evolutionary sequence, further dissection of regulatory mechanisms and their evolution during the melanophore–ML transition should provide a valuable model for understanding cell type diversification on both developmental and evolutionary time scales. The mechanisms documented here further highlight the role of positional information in generating a pigmentary phenotype, even in a species that provides a premier example of self-organization by pigment cells during body stripe formation (4, 5). Indeed, the relative importance of positional information and self-organizing processes across pattern elements and species remains an open question worthy of further study.

## Materials and Methods

### Fish Rearing and Handling

Zebrafish and other *Danio* species were reared at ∼28 °C (14L:10D) and fed rotifers, Artemia and GEMMA Micro (Skretting). Fish were anesthetized in MS222 and in most instances treated with epinephrine to contract pigment granules towards cell centers prior to imaging. Protocols were approved by the Animal Care and Use Committee of the University of Virginia.

### Mutagenesis and Transgenesis

Mutations were induced by CRISPR/Cas9 using IDT AltR reagents and analyzed after isolation by standard genetic methods or as F0 mosaics (*foxd3*). Transgenes were constructed by Gateway cloning or isothermal single reaction assembly.

### Ultrastructural and Chemical Analysis

Cytological characteristics were assessed by focused ion beam scanning electron microscopy and transmission electron microscopy. Verification of guanine association with ML used targeted and untargeted metabolomic analyses that gave equivalent results with only the latter presented here.

### Single Cell Transcriptomics and *in situ* Hybridization

Cells were collected by fluorescence activated cell sorting and captured by Chromium controller (10X Genomics) for scRNA-Seq libraries, mapped with 10X CellRanger and analyzed with Monocle3. Spatial assessment of transcript distributions followed (65).

## ACKNOWLEDGEMENTS

We thank members of the Parichy lab for assistance with fish rearing, Alishba Amjad for assistance with *in situ* hybridization, and Sarah Wilmsen and the UVA Molecular Electron Microscopy Core for assistance with electron microscopy. Supported by NIH R35 GM122471 (to D.M.P.).

## Supplementary Information Text

**Figures S1–S12**

**Movie S1–S4 legends**

**Materials and Methods**

**Tables S1–S16**

**Figure S1.**
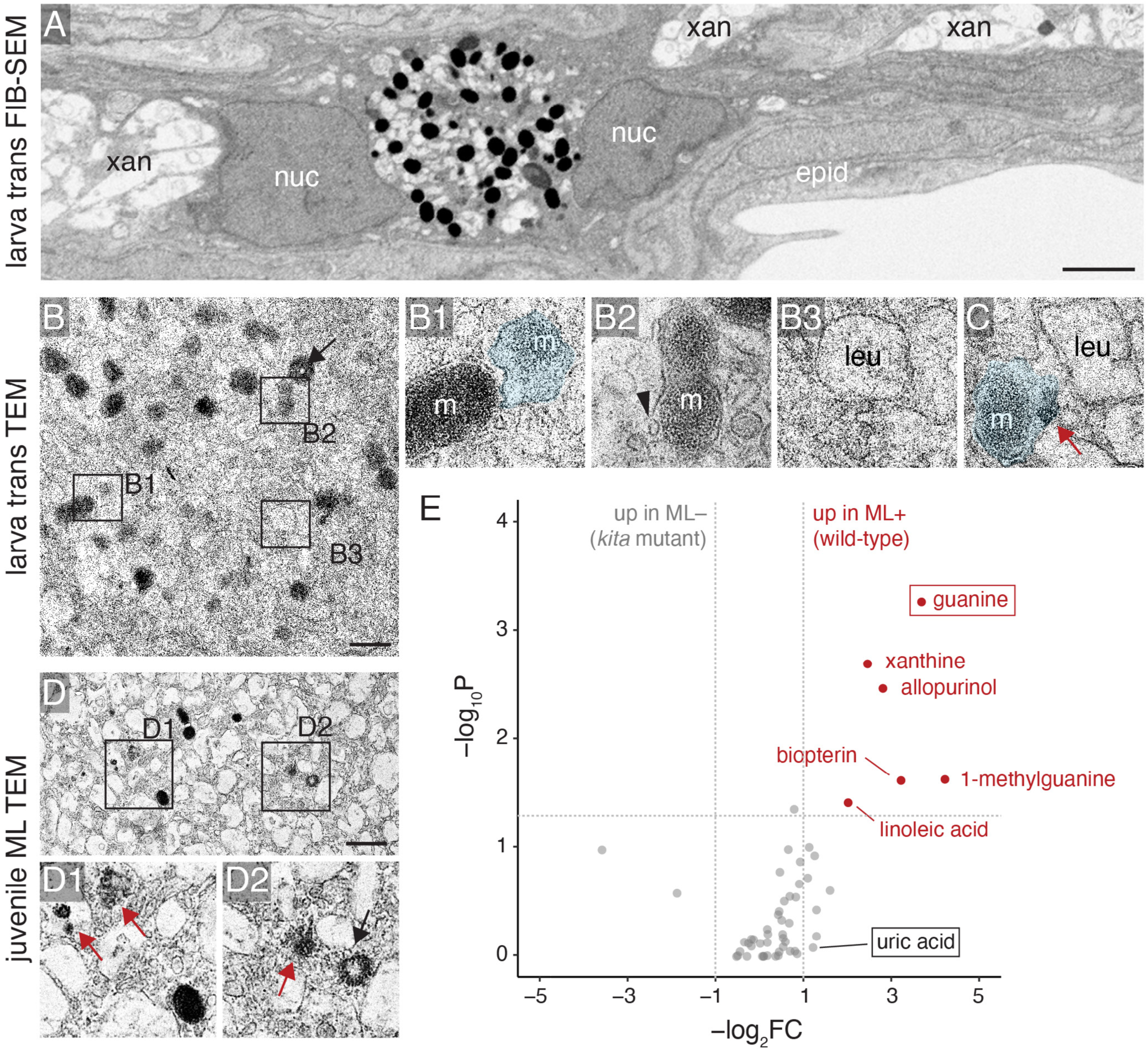
Transitional cell ultrastructure and ML chemistry. (*A*) Tissue context of cell shown in Fig. 1D, illustrating binucleation (nuc), common for ML and melanophores (1, 2), as well as nearby xanthophores (xan) and epidermal cells (epid). Melanosomes and leucosomes are concentrated in cell bodies by epinephrine treatment. (*B*) Pigmentary organelles of transitional cell, with details shown in panels *B1–3* and (*C*) from another cell at same magnification, showing irregular melanosomes (m) in various states of degradation, some associated with endosomes (arrowheads), intraluminal vesicles (black arrow), melanosome debris (red arrows) or autophagosome-like structures (blue overlays). In some cases these resembled morphologies of melanosome degradation in other systems (3). Also shown are presumptive nascent leucosomes (leu). (*D*) ML of juvenile fish contained small numbers of irregular melanosomes with leucosomes increasingly polygonal in shape. (*E*) Metabolomic analysis (*SI* Table S1) in which fins of wild-type fish with ML were compared to fins of *kita* mutants that lack ML (4). Guanine contents differed most significantly between backgrounds consistent with prior analyses of individual ML (4). Differences in pteridines may result from changes in the distribution of xanthophores, which occur in both genetic backgrounds but are associated closely with ML in the wild type. Though leucophores of medaka fish contain principally uric acid (5), an excess of uric acid was not evident in ML-containing tissue of zebrafish by this or other methods (4). These untargeted analyses further suggest that ML are unlikely to contain large quantities of previously unpredicted materials. (Scale bars: *A* 2 µm, *B* 0.1 µm, *D* 0.2 µm).

**Figure S2.**
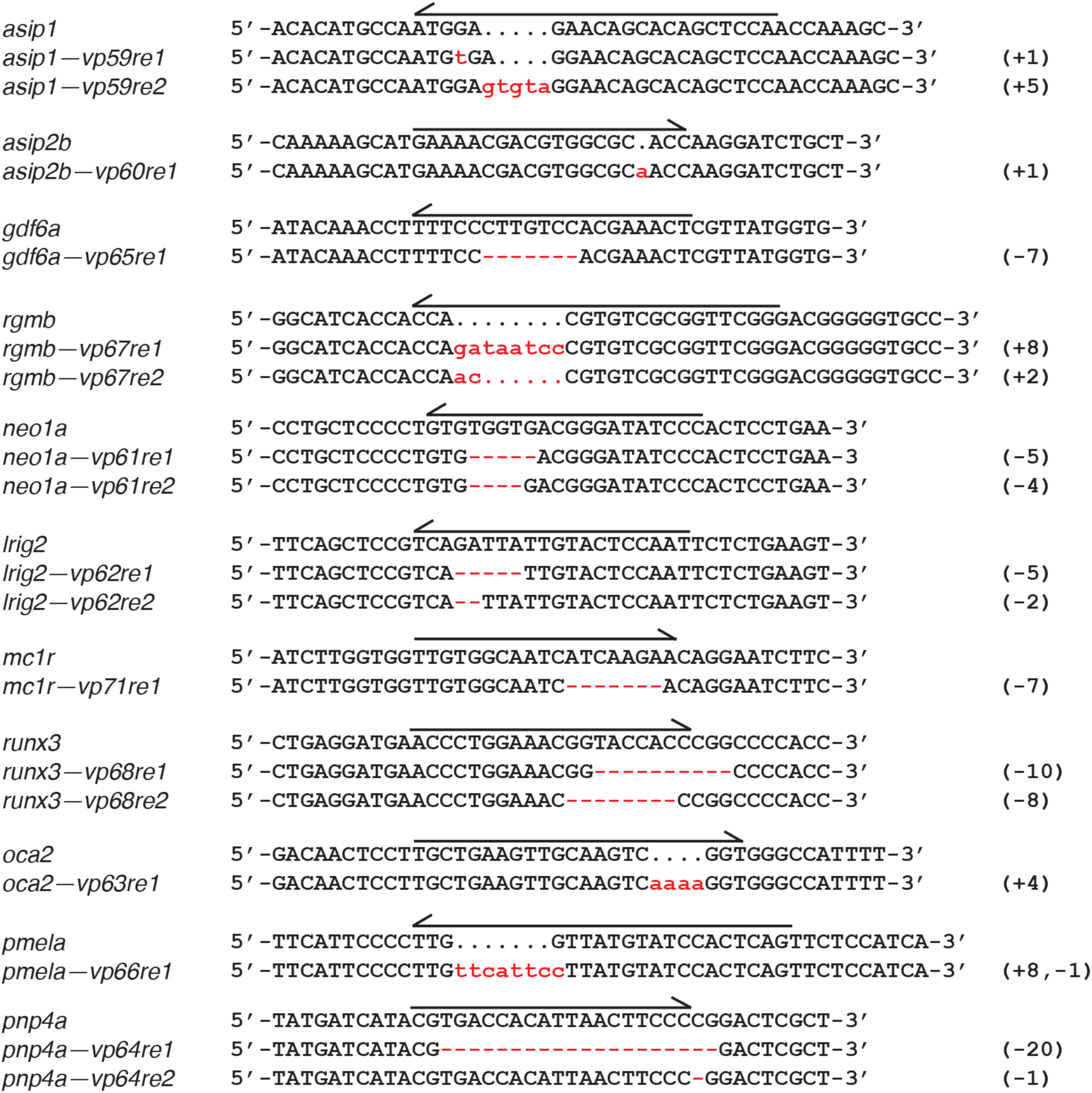
Mutant alleles. All mutations led to frameshifts and premature stop codons; differences in phenotype were not evident across alleles of each locus. Arrows denote locations of CRISPR target sites in wild-type; numbers at right, base pairs inserted or deleted.

**Figure S3.**
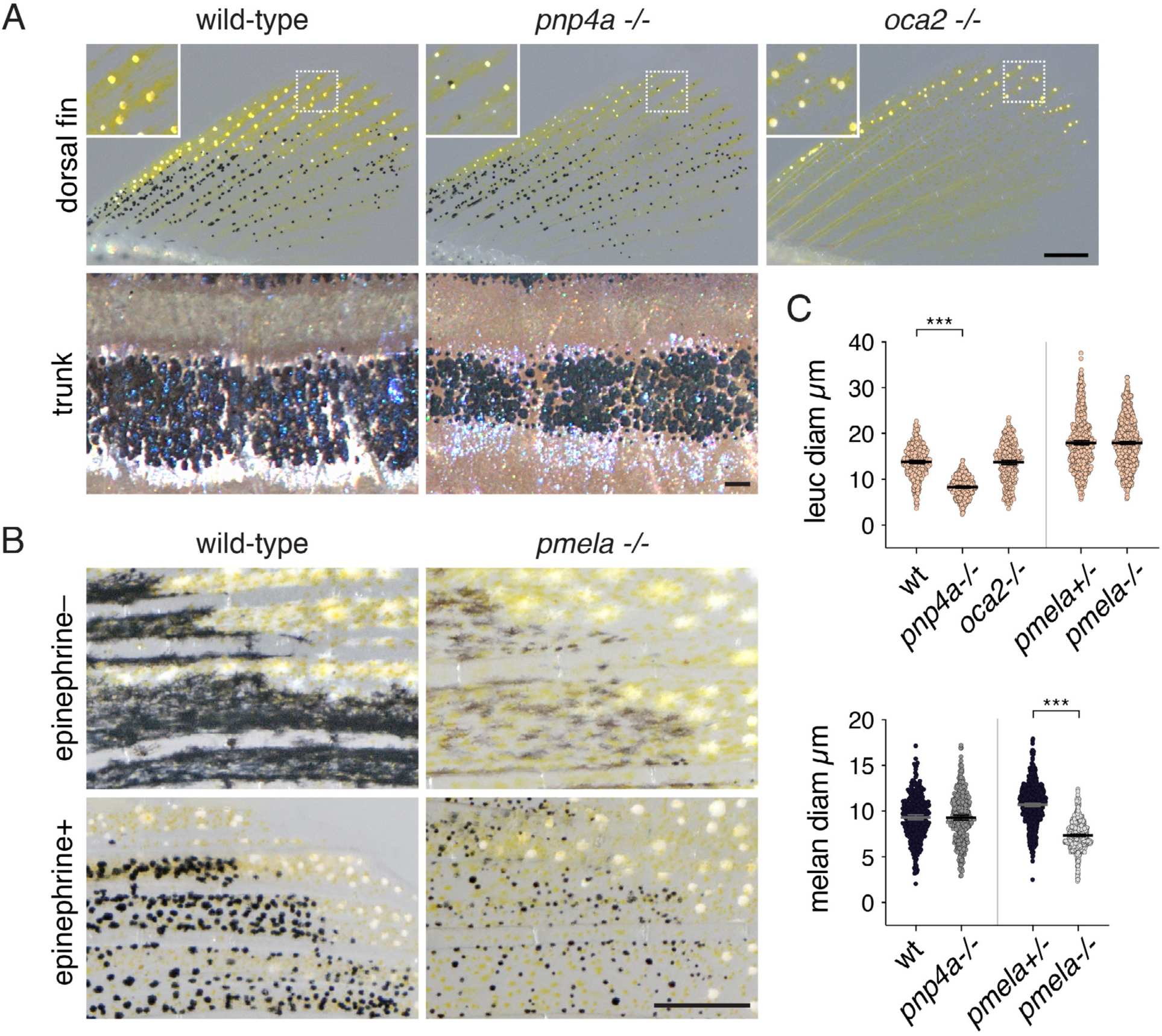
Independent pigmentary pathways of melanophores and ML. (*A*) In the fin, mutants for *pnp4a* had smaller aggregates of leucosomes following epinephrine treatment than wild-type, and on the body, they had less iridescent iridophores. (*B*) Mutants for *pmela* had less melanin than wild-type but no defects in leucophore guanine accumulation, despite moderate expression of *pnp4a* by these cells (1). Images also illustrate effects of epinephrine on both cell types, with dispersed pigmentary organelles above and contracted towards cell centers below. (C) Diameters of contracted pigment granule aggregations, indicative of total guanine or melanin accumulated (1, 6). *** Nested ANOVAs (individuals within genotypes; data *ln*-transformed to normalize residuals): leucosomes, *F*_1,9.9_=122.7, *P*<0.0001; melanosomes, *F_1,10.8_*=25.8, *P*=0.0004). *oca2* mutants completely lacked melanin and so are not shown in the plot. Total analysis: 4481 cells in 29 fish of 5 genotypes). (Scale bars: *A,B* 200 µM).

**Figure S4.**
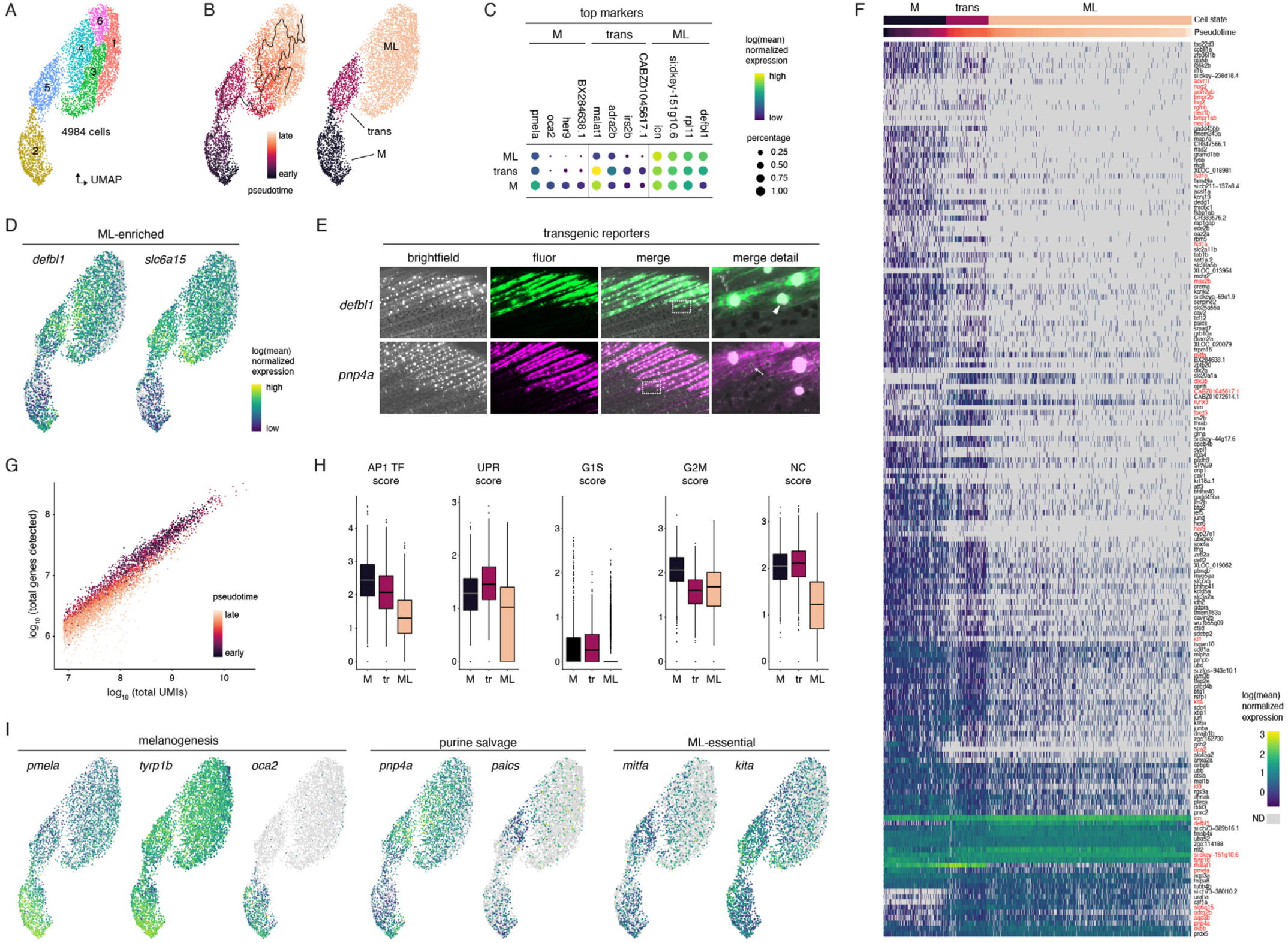
Transcriptomic state differences across melanophore-to-ML transition. (*A*) Cell groups in UMAP space identified by unsupervised clustering. (*B*) Pseudotime scores (left) from trajectory analyses with origin rooted in melanophores and assigned cell states (right) representing melanophores (M), transitional cells (trans) and ML. (*C*) Dot plots of genes identified in top marker analyses for melanophores, transitional cells and ML, showing 4 most significant hits for each state, from left to right plotted against all three cell states, top to bottom. ML mostly lacked unique markers distinguishing them from other cells states (rightmost four loci). The most abundant marker of transitional cells was *malat1* (CR383676.1), a broadly expressed long non-coding RNA that regulates gene expression at transcriptional and post-transcriptional levels (7). (*D*) Transcript abundance of two genes expressed highly in ML. The antimicrobial peptide gene *defensin beta-like 1* (*defbl1*)(8) was among the most highly expressed genes of ML compared to melanophores (log_2_FC=4.8, *q*=0.0; *SI* Table S3). Defensins can have roles in pigmentation as ligands for Melanocortin receptors (9, 10), yet fish mosaic for CRISPR/Cas9 mutations in *defbl1* did not have overt defects in ML development (*SI* Table S16). A second example locus expressed abundantly in ML, *slc6a15*, encodes a transporter of neutral amino acids (log_2_FC=2.0, *q*=4.9E-244). (*E*) Supporting the polarity of assigned state changes, differences in defbl1 expression were recapitulated by a transgenic reporter line, *Tg(defbl1:Eos)^vp57rtTg^*, with strong expression in ML (arrowhead, aggregated melanoleucosomes) and no detectable expression in melanophores. *pnp4a* transcripts were also more abundant in ML (log_2_FC=0.6, *q*=3.8E-24, panel *I*) and a second transcriptional reporter, *Tg(pnp4a:PALM-mCherry)^wprt10Tg^*, had strong expression in ML and only weak expression in melanophores (arrow). (*F*) Heatmap of gene expression plotted for cells ordered by pseudotime (top). Shown are 170 genes selected for having the most significant expression differences by p-value across cell states (M, trans, ML), or because they were highlighted as informative, differentially expressed or functionally required in other analyses (genes in red; see main text and following figures). Transcripts of most loci were less abundant in ML as compared to melanophores or transitional cells. (*G*) Total genes detected increased with numbers of unique molecular identifiers (UMIs), representing depth of sequencing per cell. Yet detected genes were also lower for cells with later pseudotime scores, for a given UMI, implying that fewer genes are expressed as cells transition to the ML state. (*H*) Box plots indicate differences in transcriptomic signature scores across cell states (gene sets provided in *SI* Table S5). AP1 transcription factor gene expression is associated with overall transcriptional activity whereas unfolded protein response (UPR) gene expression is associated with overall translational activity (1). Relatively higher scores for G2–M cell cycle genes across M–ML as compared to G1–S cell cycle genes may reflect a terminal, acytokinetic mitosis in which many of these cells become binucleated (e.g., *SI Appendix* Figs. *S1A*) and proliferatively arrested, as is true of melanophores on the body (1). Transitional cells (tr) did not have a markedly higher signature score for genes of embryonic neural crest (NC) cells suggesting that M–ML transition does not entail a generalized dedifferentiation, as observed for some cell types during regeneration (11) and in some melanoma cells (12). (*I*) Transcript abundance for selected loci mapped onto cells in UMAP space, including genes of the melanin and purine synthesis pathways, and genes previously identified to be essential during the transition of melanophores to ML (4). (Scale bar: *F* 200 µm).

**Figure S5.**
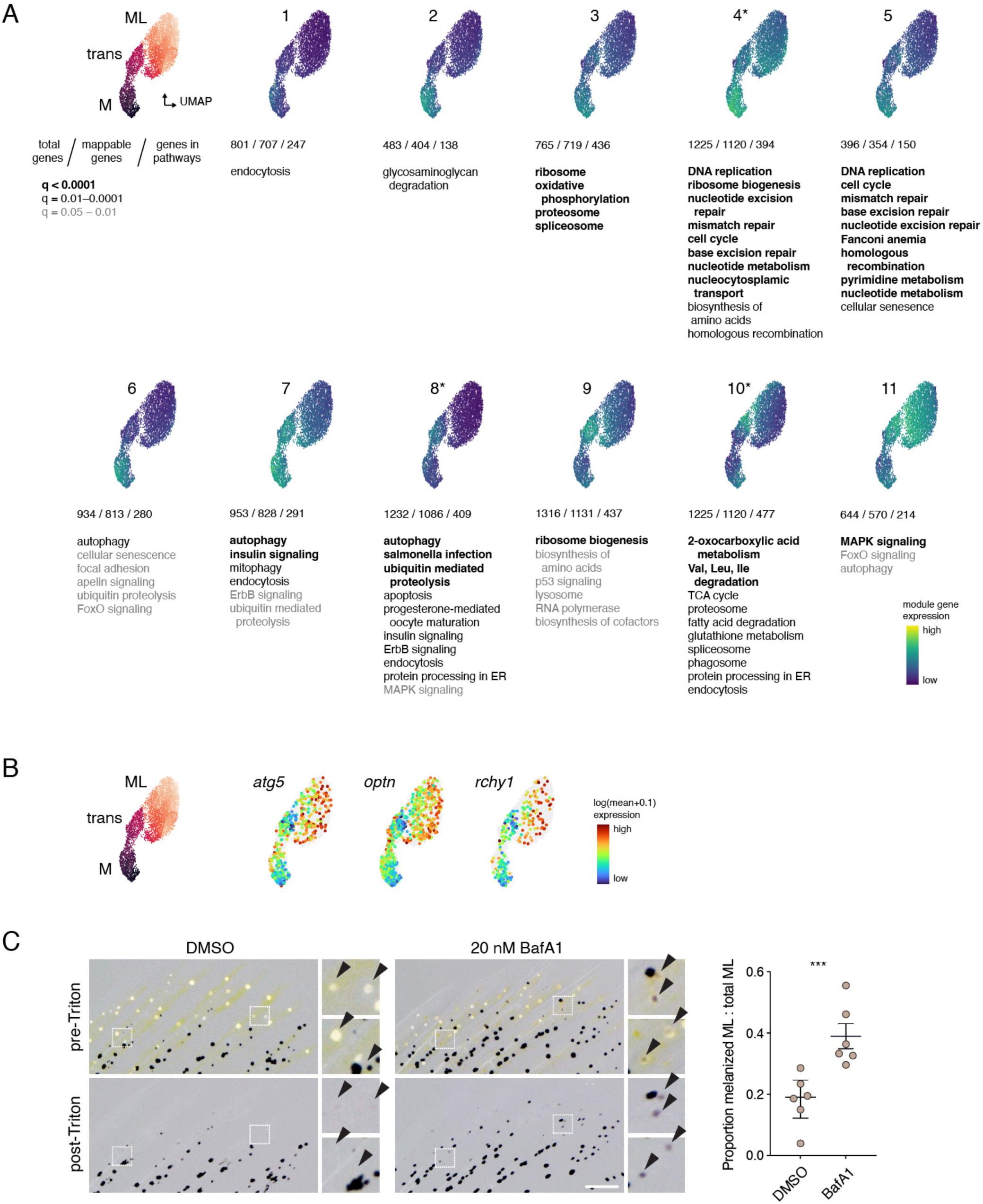
Coexpressed modules of genes across melanophore-to-ML transition. (*A*) Gene module expression mapped onto cells in UMAP space, ordered in approximate sequence with differentiation towards ML. Modules were identified in Monocle 3 (13, 14) as sets of genes covarying with one another and with pseudotime scores, or sets of genes covarying independent of pseudotime score (marked by *). Beneath each plot are total genes identified in each module, genes mappable to orthologous loci in KEGG pathways, and genes in KEGG pathways having associations of *q*<0.05. Pathways are indicated with different type faces representing strengths of association (by *q* values). (*B*) In transitional cells, ultrastructural features of melanosomes were consistent with degradative mechanisms and autophagy pathway genes appeared in modules 7 and 8* of panel A. In mammals, orthologues of *atg5*, *optn* and *rchy1* play important roles in melanosome autophagy (15). Average transcript abundances for these loci did not differ across cell states, yet patterns of expression were altered, with many melanophores expressing each locus at low levels but a small proportion of ML expressing each locus at much higher levels. Cells with detectable transcripts are shown with larger points to aid visualization, despite sparsity overall. (*C*) Fins of fish treated with DMSO or bafilomycin A1 that were fixed with 4% paraformaldehyde and then rinsed within Triton X-100 to dissolve guanine from ML. Insets show ML before and after guanine removal, with persisting melanin present in bafilomycin A1 treated ML. Detergent treatment also removed yellow pigmentation from xanthophores. Plot shows proportions of ML with persisting melanin relative to the total population of ML within each fin (*n*=12 fish; likelihood ratio test, χ^2^=13.0, d.f.=1, *P*<0.0003). Lower images have been warped manually in Photoshop to align corresponding cell positions before and after Triton X-100 treatment (e.g., arrowheads). (Scale bar: *C* 100 µm)

**Figure S6.**
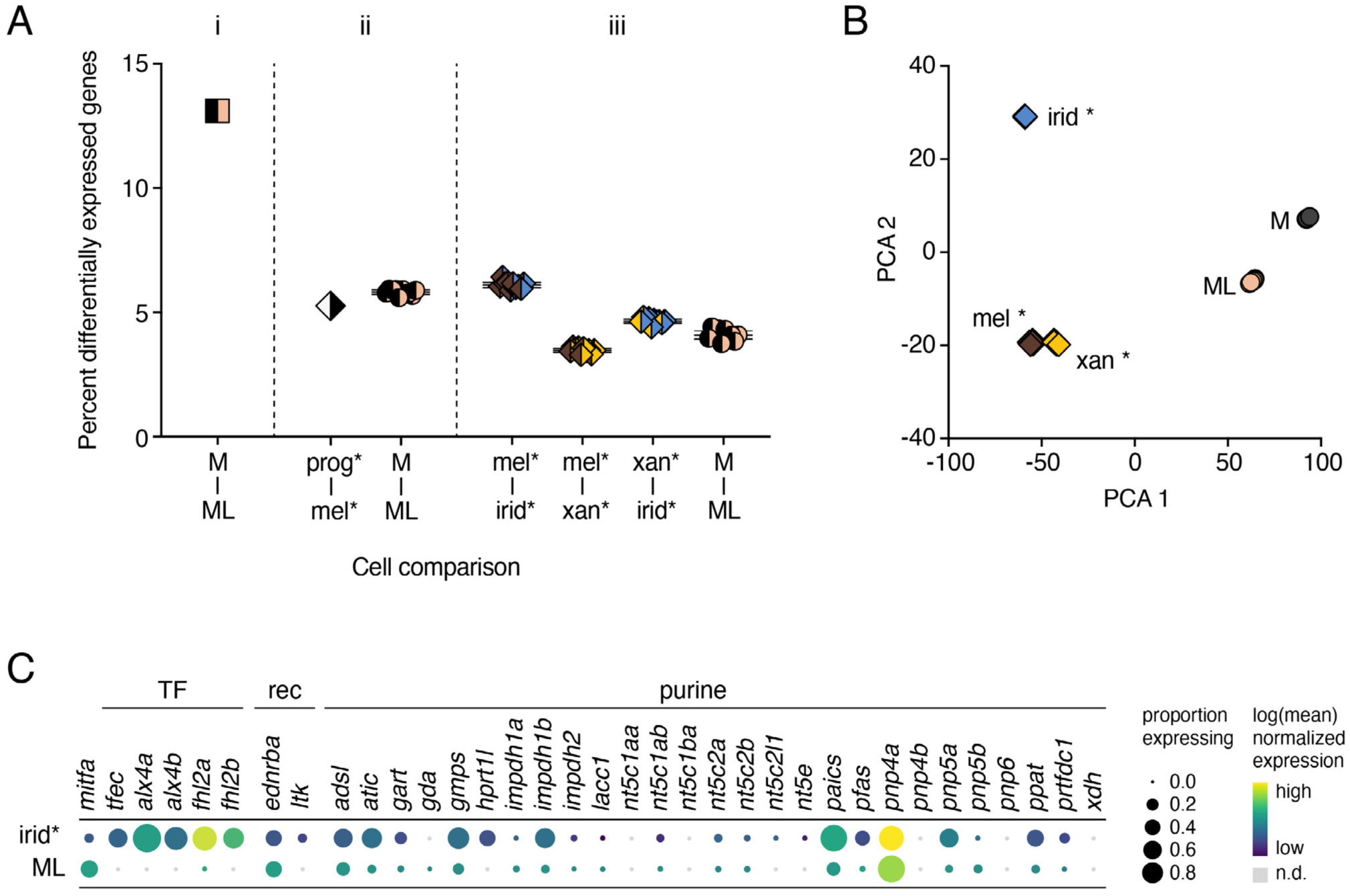
Transcriptomic state differences among pigment cells and progenitors. (*A*) Melanophore and ML transcriptomic state differences were of similar magnitude to those of other pigment cell classes and progenitor–derived states. Plot shows differentially expressed genes (DEG) as percentage of total genes detected for comparisons indicated on X-axis: melanophores and ML on the fin in this study (M–ML) and cell types (*) identified in a comparable study of pigment cell development on the body (mel, melanophore; xan, xanthophore; irid, iridophore; prog, multipotent pigment cell progenitors) (1). Genes were considered differentially expressed when log_2_FC>1 and *q*<0.05. *Plot segment i*, Comparison of M and ML of the fin identified 13.1% of detected genes as differentially expressed (M, 1008 cells; ML, 3026 cells). *Plot segment ii*, Comparison of progenitor and derived cell types yielded similar percentages of differentially expressed genes for the body (prog–mel, 390 and 638 cells, respectively) and the fin (M–ML, downsampled randomly to 390 cells and 638 cells, respectively, with 10 replicate comparisons). Bars, mean±95% CI. *Plot segment iii*, Comparisons of differentiated cells. On the body, melanophores and iridophores (mel–irid) had the most differentially expressed genes, whereas M–ML differences were on par with those of other comparisons (mel–xan; xan–irid; all downsampled randomly to 400 cells of each type, with 10 replicate comparisons each). (*B*) Averaged states of pigment cell transcriptomes in PCA space show separation of M and ML greater than observed for melanophores and xanthophores, but not as extreme as for comparisons of iridophores to melanophores or xanthophores. The difference in multivariate space between iridophores and ML is particularly notable given the phenotypic dependence of both cell types on crystalline guanine. Points represent each of 10 replicate downsampled datasets comprising 400 cells each, treated as pseudobulk RNA-Seq data for PCA analysis. (*C*) Dot plot highlighting differences in gene expression profiles between iridophores (irid*) and ML, with transcription factors of iridophores (TF; *mitfa* shown for comparison), receptors essential for iridophore development (rec), and genes of the purine synthesis pathway (1, 16–23). Differences in transcription factor expression suggest that ML and iridophores have distinct gene regulatory programs upstream of guanine synthesis and crystal formation.

**Figure S7.**
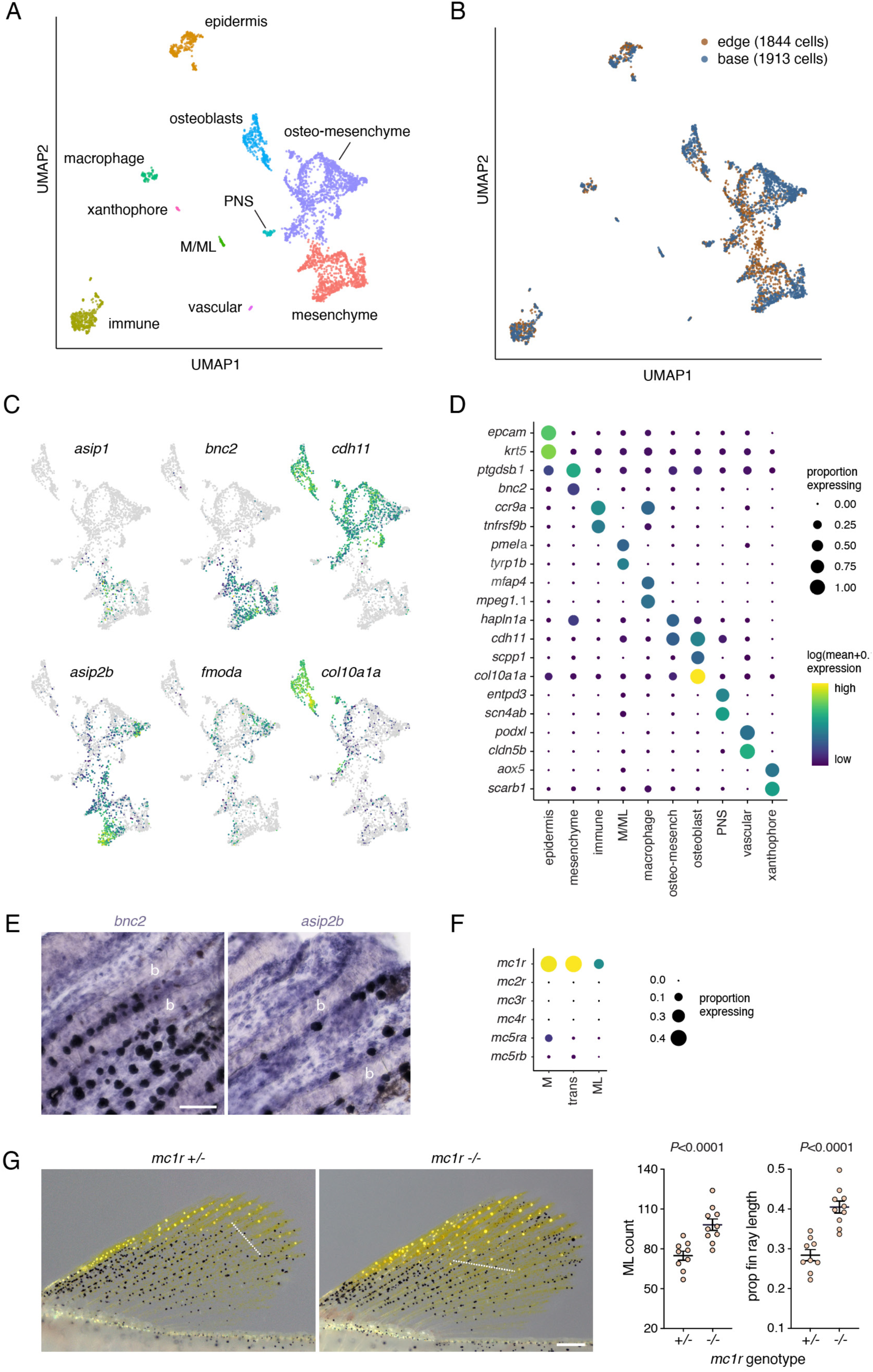
Comparison of fin edge and base and roles for Agouti signaling. (*A,B*) Single cell transcriptomes in UMAP space showing cell-type assignments and complements of cells from the anterior/distal fin edge vs. fin base (including ventral/posterior edge; locations in Fig. 2D). (*C*) Transcript abundances mapped onto cells in UMAP space for *asip1* and *asip2b* as well as markers of non-osteoblast fin mesenchyme (*bnc2*), osteo-mesenchymal cells (*fmoda*), osteo-mesenchymal cells and osteoblasts (*cdh11*), and osteoblasts (*col10a1a*) (Fig. 2B). (*D*) Top markers of assigned cell types. (*E*) *In situ* hybridization for *bnc2* and *asip2b* transcripts, neither of which differed in abundance between edge and base. b, bones of fin rays. Images are projections of multiple focal planes. (*F*) Melanophores– ML expressed melanocortin receptor genes *mc1r*, *mc5ra* and *mc5rb* at detectable levels. (*G*) Mutation of *mc1r* led to more ML that extended further towards the fin base than in wild-type. Dashed lines indicate the averaged positions across the most basal ML of 3^rd^, 4^th^ and 5^th^ fin rays, used in calculating average proportions of fin rays covered by ML at right (*SI* Table S10). (Scale bars: *E* 50 µm, *G* 200 µm).

**Figure S8.**
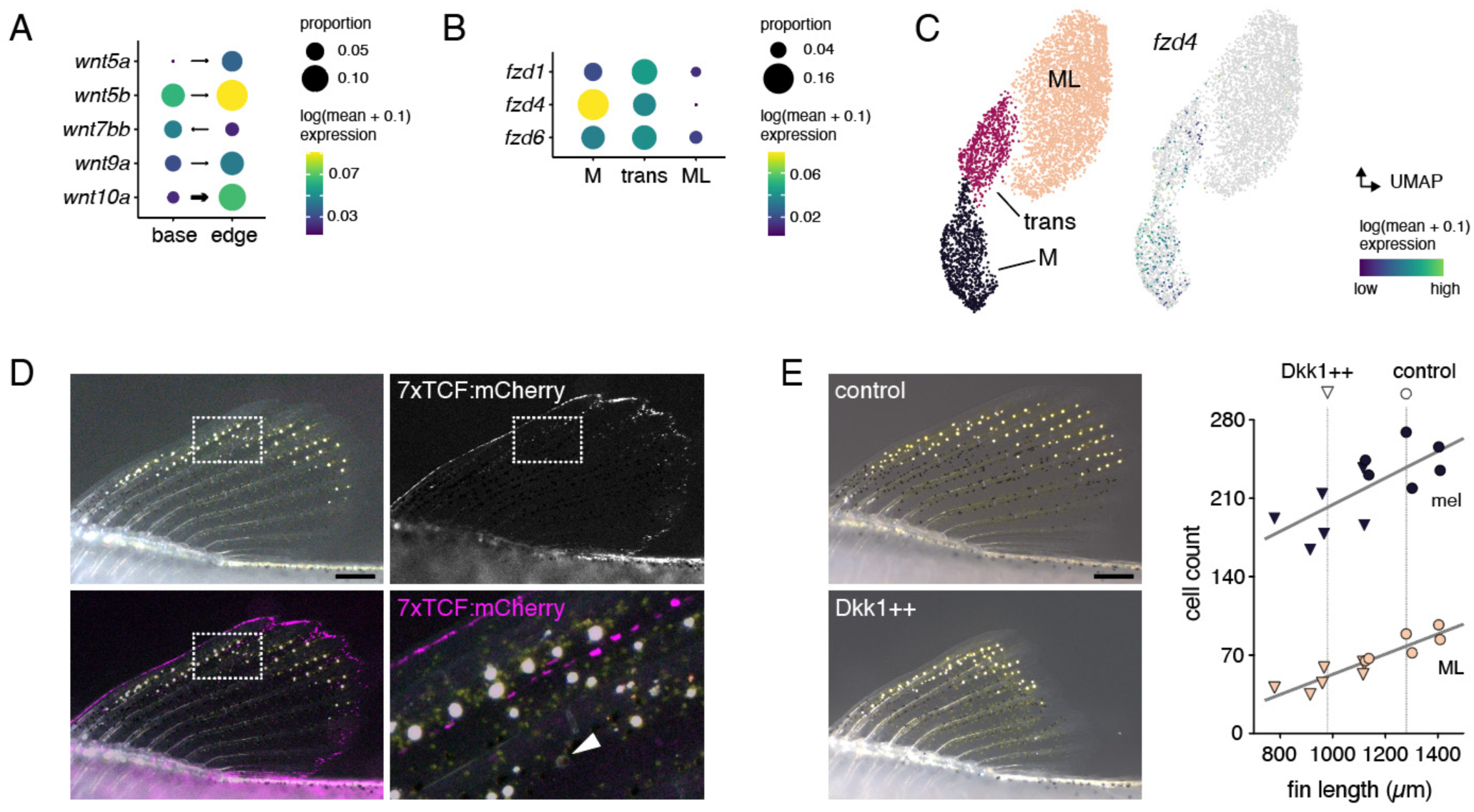
Wnt signaling affected fin size but not ML development. (*A*) Expression of genes encoding ligands of Wnt pathways in the tissue environment of melanophores and ML at fin base and edge, showing proportion of cells with detectable transcripts and mean abundances of transcripts in those cells. Arrows point to fin region with significantly higher transcript abundance in tests of differential expression (thin arrows: *p*<0.05, *q*>0.05; wide arrows: *p*<0.05 and *q*<0.05), with sampled cells corresponding to those of *SI Appendix* Fig. S7C and Table S8. (*B*) Expression of Wnt pathway receptor genes, filtered for loci expressed by >10% of cells in at least one cell state and having mean abundances of >1 transcript per expressing cell. Values for these and additional loci are provided in *SI* Table S11. (*C*) Expression of *fzd4* mapped onto cells in UMAP space. (*D*) Wnt signaling reporter of *Tg(7xtcf-xla-siam:mCherry)^ia4Tg^* (24) was detectable in cells along the fin edge and in sparsely distributed cells more centrally, but was not evident in ML or transitional cells (arrowhead in detail of bright field and fluorescence merged image, lower right) despite contraction of pigment granules to cell centers by epinephrine treatment. (*E*) Wnt pathway inhibition by repeated post-embryonic heat shock of *hs:dkk1* (25, 26) blocked fin outgrowth as measured by length of the third fin ray (*F*_1,10_=16.9, *P*=0.002); vertical dashed lines in plot indicate mean sizes of control fins (circles) and Dkk1-overexpressing fins (triangles). Analyses of covariance showed significant increases in melanophore and ML numbers with fin size (regression coefficients±SE: 0.12±0.04, *P*<0.01 and 0.09±0.01, *P*<0.0001, respectively), but no differences in cell numbers due to Dkk1-overexpression after controlling for effects on fin size (*F*_1,9_=2.11, *P*=0.2 and *F*_1,9_=1.08, *P*=0.3, respectively).

**Figure S9.**
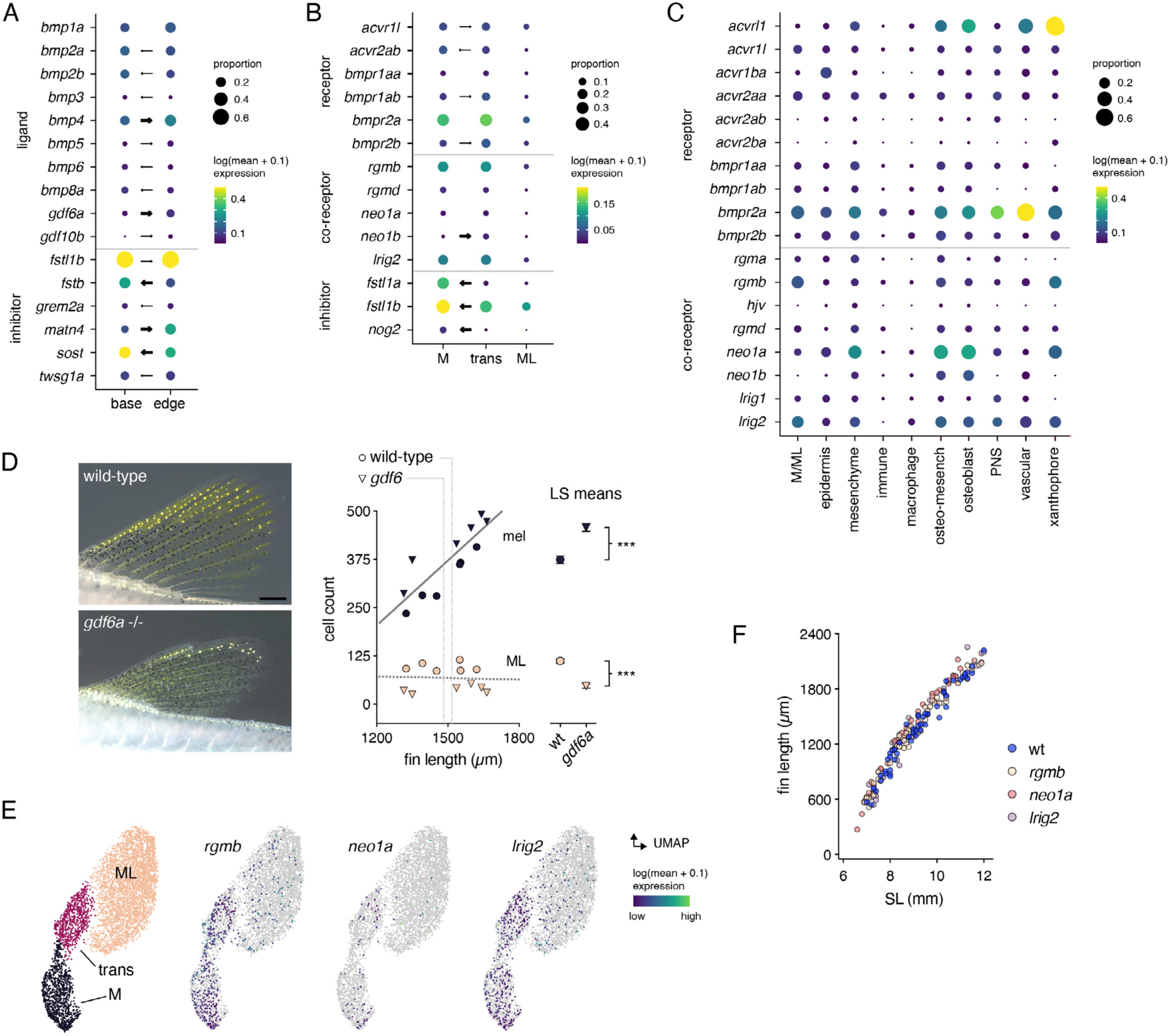
BMP pathway gene expression and dependence of ML on BMP ligand Gdf6a. (*A*) BMP ligand genes expressed in the fin tissue environment (as in *Supplementary Appendix* Fig. S7A). Of ligand genes having transcripts with greater abundance at the fin edge, *gdf6a* had the largest difference in expression relative to the fin base (log_2_FC=1.18, *q*=3.46E-05). *bmp4* was expressed more robustly and was detectable in more cells yet had a smaller difference in transcript abundance between edge and base (log_2_FC=0.62, *q*=0.005). Also shown are 6 genes that encode secreted inhibitors of BMP activity (out of 18 such genes detected) that also exhibited significant differences in transcript abundance between regions. Arrows point to region with higher expression (thin arrows: *p*<0.05, *q*>0.05; wide arrows: *p*<0.05 and *q*<0.05). (*B*) Genes encoding BMP/TGFβ receptors, co-receptors and secreted inhibitors expressed by melanophores–ML. (*C*) BMP/TGFβ pathway receptor and co-receptor gene expression in fin tissue environment by cell type. BMP coreceptor *rgmb* was expressed by a relatively high proportion of the few M/ML (*n*=41) captured (*SI Appendix* Fig. S7A) as compared to other cell types (total cells=3757). For *A–C*, genes are shown only if expressed by at least 5% of cells in a category. Additional BMP and TGFβ pathway receptors are listed in *SI* Table S11. (*D*) Mutants for *gdf6a* had fewer ML than wild-type siblings. Among fish selected to have similar fin sizes, *gdf6a* mutants had more melanophores and fewer ML than wild-type siblings, with results for melanophores consistent with published observations (27). Melanophore but not ML numbers varied with fin size (regression coefficients±SE: 0.53±0.05, *P*<0.0001 and 0.01±0.03, *P*=0.6, respectively). Least squares mean±SE numbers of melanophores and ML after controlling for fin length are shown at right (*F*_1,8_=34.51, *P*<0.0001 and *F*_1,8_=73.53, *P*<0.0001, respectively). (*E*) *rgmb*, *neo1a* and *lrig2* expression by melanophore–ML mapped in UMAP space. (*F*) Mutants for *rgmb*, *neo1a* and *lrig2* did not have fin lengths significantly less than wild-type (Dunnett’s one-tailed d=2.09, all mutant genotypes *P*>0.8, after controlling for overall covariance of fin length and standard length, *P*<0.0001). (Scale bar: *D* 200 µm).

**Figure S10.**
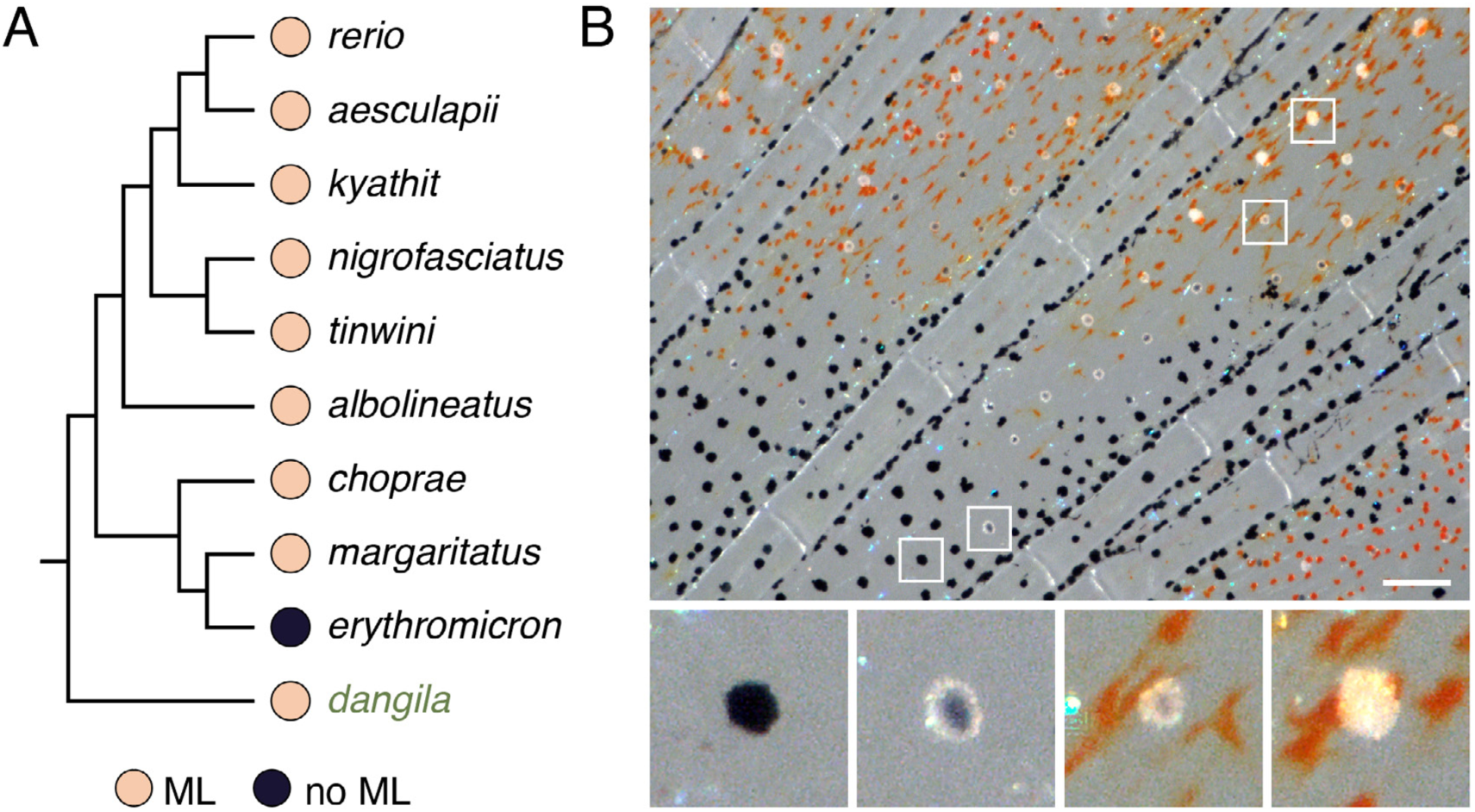
Melanoleucophores occur in multiple *Danio* species. (*A*) Nine of 10 previously examined species (in black) had ML in their dorsal fins (4); phylogeny according to (28). (*B*) The large-bodied species *D. dangila* is positioned basally in the phylogeny, co-occurs with *D. rerio* (29), and is shown here to have ML as well, with different degrees of residual melanin shown. Though species of genera most closely related to *Danio* seem not to have melanoleucophores (e.g., *Devario*, *Chela*)(30, 31) it remains uncertain whether these cells are uniquely shared and derived within *Danio* or whether convergently evolved (or re-evolved) melanoleocophores might contribute to white fin ornamentation evident in some more distantly related taxa. (Scale bar: *B* 100 µm).

**Figure S11.**
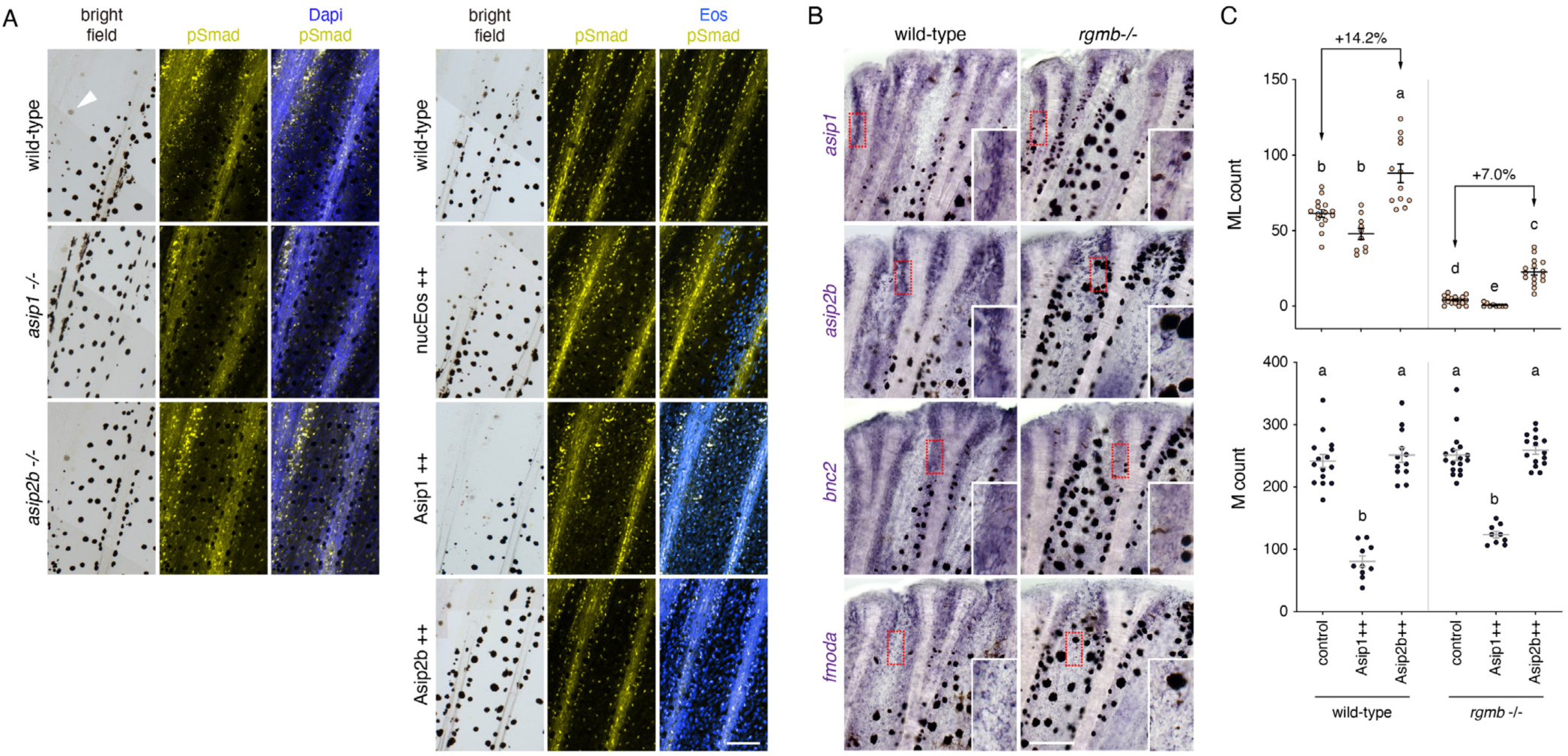
BMP and Agouti pathway dependencies. (*A*) pSmad staining for BMP signaling activity did not differ from wild-type in fish mutant for *asip1* or *asip2b* (left), or overexpressing Asip1 and Asip2b mosaically (right). nucEos, fluorophore-only overexpression control pseudocolored blue. Arrowhead, transitional cell (transmitted illumination). (*B*) *In situ* hybridization in wild-type and *rgmb* mutant fins. Staining for *asip1*, *asip2b*, *bnc2* and *fmoda* were somewhat diminished in the mutant. Insets, higher magnification of regions outlined in red. (*C*) Effects of Asip1 and Asip2b overexpression in wild-type vs. *rgmb* mutant siblings for ML and melanophores. *rgmb* homozygosity reduced ML counts compared to wild-type overall, whereas Asip1 further lowered ML numbers and Asip2b increased numbers (as in Fig. 2G). Nevertheless, the effects of Asip2b dependent on background with a more substantial increase in wild-type than *rgmb* mutants (genotype x transgene effect interaction, *F*_1,55_=7.1, *P*=0.006). Interactions of this sort were not evident for effects of Asip1 on ML, nor for either transgene on melanophores. Shared letters indicate groups not significantly different from one another in *post hoc* Tukey Kramer comparison of means. Counts for ML were squareroot transformed for analyses of variance to normalize variance in residuals across groups.

**Figure S12.**
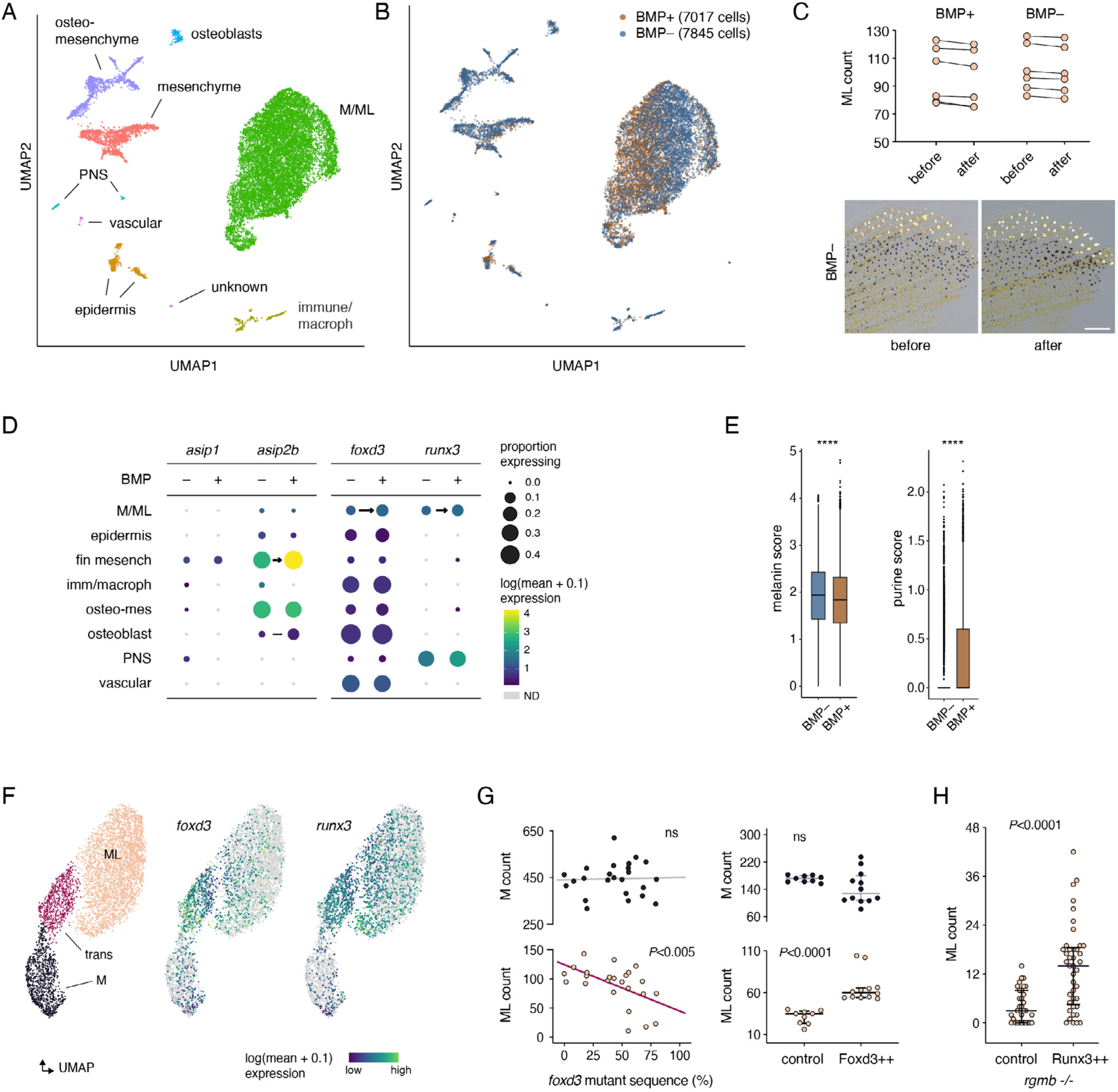
Comparison of single cell transcriptomes in response to BMP signaling inhibition. (*A*) Identified cell types in UMAP space. (*B*) Vehicle treated control cells (BMP+) and LDN-193189 treated cells (BMP–). (*C*) Upper, ML numbers before and after treatment did not differ between BMP+ and BMP– groups (matched pairs, *F*_1,10_=0.01, *P*=0.9; *SI* Table S12). (*D*) Dotplots for Agouti genes as well as *foxd3* and *runx3* expression in response to BMP-inhibitor treatment. Arrows indicate differences (where *q*<0.05) and condition of higher transcript abundance in gene expression comparisons of BMP inhibitor effects within cell types indicated. BMP inhibitor treatment did not affect abundance of melanocortin receptor transcripts in M–ML (*mc1r*, *mc5ra*: *q*=0.9, 0.2, respectively; not included in the plot). (*E*) Signature scores for genes of melanin and purine synthesis/salvage pathways reveal statistically significant differences (*P*<0.0001). (*F*) In unmanipulated cells, transcripts for *foxd3* and *runx3* were expressed more abundantly in transitional cells than either melanophores or ML (*foxd3*: log_2_FC=1.7, *q*=1.15E-122; *runx3*: log_2_FC=1.3, *q*= 1.18E-97; *SI* Table 4). *core-binding factor subunit beta* (*cbfb*) encoding a primary Runx3 heterodimerization partner promoting DNA binding affinity and stability (32, 33), was also upregulated in transitional cells (log_2_FC=0.4, *q*= 1.39E-05). Low levels of *runx2b*, but not other Runx genes, were also evident in M–ML. (*G*) Left, Fish mosaic for *foxd3* mutations had reduced numbers of ML as mutagenesis efficiency in sampled fin tissue increased. Right, Heat-shock induced Foxd3 overexpression (Foxd3++) increased ML complements (Wilcoxon *S*=45, *P*<0.0001; bars, median±interquartile range; *SI* Table S15). (*H*) Mosaic heat-shock induced Runx3 (Runx3++) increased ML complements of *rgmb*−/− mutants (Wilcoxon *S*=805, *P*<0.0001). Scale bar: *C* 200 µm).

## SI Movies

**SI Movie S1.** FIB-SEM imaging of melanophore during stage of melanophore–ML transition. Pigments were contracted to cell centers by treating with epinephrine. Two melanophores with homogeneous melanosomes, adjacent to xanthophores that contain pterinosomes (white) and carotenoid vesicles (dark). Scale bar, 5 µm.

**SI Movie S2**. FIB-SEM imaging of transitional cell illustrating abnormal melanosomes and early leucosomes. The transitional cell at the center was surrounded by xanthophores, with white pterinosomes and dark carotenoid vesicles. Scale bar, 5 µm.

**SI Movie S3**. FIB-SEM imaging of adult melanophore with hundreds of homogeneous melanosomes. Scale bar, 5 µm

**SI Movie S4.** FIB-SEM imaging of adult melanoleucophore showing large and jagged leucosomes, with very sparse melanosomes or debris. Scale bar, 5 µm.

## Materials and Methods

### Fish Stocks and Rearing Conditions

Fish were reared under standard conditions (∼28 °C; 14L:10D) with larvae fed initially marine rotifers, derived from high-density cultures and enriched with Algamac (Reed Mariculture). Older larvae and adults were transitioned to live brine shrimp and GEMMA Micro 500 (Skretting). Stocks of *D. rerio* were: *asip1^vp59re1^, asip1^vp59re2^*, *asip2b^vp60re1^*, *gdf6za^vp65re1^*, *kita^b5^* (34), *lrig2^vp62re1^*, *lrig2^vp62re2^*, *mc1r^vp71re1^*, *neo1a^vp61re1^*, *neo1a^vp61re2^*, *pmela^vp66re1^*, *pnp4a^vp64re1^*, *pnp4a^vp64re2^*, *oca2^vp63re1^*, *rgmb^vp67re1^*, *rgmb^vp67re2^*, *runx3^vp68re1^*, *runx3^vp68re2^*, *Tg(hsp70:asip1-2a-neos)^vp72rTg^*, *Tg(hsp70:asip2b-2a-neos)^vp73rTg^*, *Tg(defbl1:EosFP)^vp57rtTg^*, *Tg(hsp70:dkk1)^w32Tg^* (25), *Tg(hsp70:foxd3-2a-nuc-mcherry)^vp72rTg^*, *Tg(pnp4a:PALM-mCherry ^wprt10Tg^* (35), *Tg(tyrp1b:PALM-mCherry)^wprt11^* (36), *Tg(7xtcf-xla-siam:mCherry)^ia4Tg^* (24). *Danio albolineatus* were derived from individuals collected in Thailand by M. McClure in 1995, provided to the laboratory of S. Johnson, and then maintained in our laboratory from 2000 until the present. *Danio dangila*, which co-occur with *D. rerio* (37), were obtained from a commercial supplier. This study was performed in accordance with the recommendations in the Guide for the Care and Use of Laboratory Animals of the National Institutes of Health. Animals were handled according to approved institutional Animal Care and Use Committee (ACUC) protocol (#4170) of the University of Virginia. Euthanasia was accomplished by overdose of MS222 (Syndel) followed by physical maceration.

### CRISPR/Cas9 Mutagenesis

CRISPR/Cas9 mutants or F0 mosaics were generated by injecting one-cell stage embryos with approximately 1 nl of 5 µM gRNA:Cas9 RNP complex (IDT)(38). For production of mutant lines, fish were sorted for dorsal fin phenotypes at juvenile stage (39)and alleles recovered by incrossing and outcrossing. For F0 mosaic analysis, injected embryos were raised to juvenile stage, then imaged and genotyped with DNA extracted from dorsal fins. Mutagenesis scores were estimated using TIDE software (<u>tide.nki.nl</u>).

### Transgenesis

*hsp70:asip1-2a-neos*, *hsp70:asip2b-2a-neos*, *hsp70:foxd3-2a-nlsmCherry* and *hsp70:runx3-2a-EGFP* plasmids were assembled using the *hsp70l* promoter and destination plasmids from the Tol2kit (40), coding sequences were cloned from skin wild-type cDNA, and linked to fluorophores or photoconvertible fluorophore EosFP with a nuclear-localizing sequencing using viral 2a sequence (41). Plasmids were injected at the one-cell stage with *tol2* mRNA at 25 pg per embryo (42). Injected embryos were sorted for fluorophore after 40 °C heat-shock for 1 hr at 3 days post fertilization. Integrated transgenes were recovered by intercrossing and outcrossing.

### Heat-shock Induction

Fish at the stage of ML initiation (∼7 mm standard length)(39) were placed in tanks on a custom built heat-shock rack to receive 1 heat-shock on day 1, followed by 6 heat-shocks per day for 2 weeks. Heat-shocks were 1 hr, using fresh fish water pre-warmed to 39 °C.

### Pharmacological Analysis

Fish were treated chronically with 2 µM LDN-193189 (S2618, Selleckchem) / 0.5% DMSO, or 20 nM Bafilomycin A1 (S1413, Selleckchem) / 0.5% DMSO starting from 7 mm SL for 2–3 weeks. Fish were released to fresh water and received brine shrimp in the day and were immersed in fresh drug solution overnight. For acute treatment, fish at 12 mm SL were treated with 2 µM LDN-193189 (S2618, Selleckchem) / 0.5% DMSO for 2 d.

### Immuno-fluorescent Antibody Staining

Fins were fixed in 4% PFA for 1 hr at room temperature, incubated in blocking solution overnight and primary antibody against pSmad (1:100 dilution; Phospho-Smad1/5/9, 13820S, Cell Signaling Technology) overnight. Samples were washed with PBST (0.1% Triton X-100 in 1x PBS) 5 times for 5 min, then incubated overnight with fluorescent dye conjugated secondary antibody before washing and imaging.

### Imaging, Documentation and Analysis

Fish were anesthetized in MS222 prior to imaging. To contract pigment granules for imaging, fish were treated with 1 mg/ml epinephrine (E4642, Sigma-Aldrich) for 5 min. Bright-field and low-resolution fluorescent images of fish fins or bodies were acquired using Zeiss AxioZoom stereomicroscope equipped with Zeiss Axiocam cameras, or Zeiss AxioObserver inverted microscope equipped with Yokogawa CSU-X1M5000 laser spinning disk and Hamatsu camera. High-resolution fluorescence images and corresponding pseudo-brightfield images were acquired using a Zeiss LSM880 inverted laser confocal microscope in Airyscan SR mode. Images were captured either as single frames or as tiled sets of larger areas that were then stitched computationally using ZEN Blue software. The same acquisition parameters were applied across matched sets of images (e.g., across genotypes and treatments) as were subsequent adjustments to color balance, display levels or both, applied to entire images as needed for visualization in Adobe Photoshop CS. Phenotype documentations were done with ZEN Blue software, including counts of pigment cell numbers, measurements of fluorescence signal intensities, and measurements of pigment aggregations within cells, and lengths of fin bases and 3rd fin rays. To reveal residual melanin in ML, guanine deposits were removed by incubation of fin tissue with 0.1% Triton X-100. Analyses of cell counts and other attributes were conducted in JMP Pro 18.0.1 for Apple Macintosh (JMP Statistical Discovery; jmp.com).

### Focused Ion Beam Scanning Electron Microscopy (FIB-SEM) and Transmission Electron Microscopy (TEM)

Fins were amputated and pre-fixed for 15–30 min with 4% formaldehyde and 2.5% glutaraldehyde in PBS, then changed for an 2 additional hr into fresh 4% formaldehyde and 2.5% glutaraldehyde in 100 mM phosphate buffer at pH 7.4 at room temperature. Tissues were post-fixed for 2 hr with 2% osmium tetroxide, 2% uranyl acetate for 2 hr at room temperature and dehydrated with acetone and flat embedded in epoxy resin. Volume microscopy was performed using a focused ion beam scanning electron microscope (Helios 650; FEI Company, Eindhoven, The Netherlands) (43). Milling was performed with a gallium ion beam at 30 kV and 2.5 nA, and the imaging was done at 1.5 keV, 800 pA, 6144 × 4096-pixel frame size, 6 µs dwell time, with 100 Å pixel size, and 200 Å section thickness (Kizilyaprak et al., 2015). After milling at 52°, the fresh surface was tilted to 90° and imaged normally to the electron beam (44). Images were aligned with IMOD (45). TEM imaging for mature ML and leucosomes was done with the same FIB-SEM sample block. After acquiring data by FIB-SEM, the sample blocks were mounted in an ultramicrotome (Leica UC7, Leica Microsystems, Vienna, Austria) and sections cut at 700–1000 Å with a diamond knife (Diatome LTD, Nidau, Switzerland). Imaging was done in a TEM (L120, Thermo Fisher Scientific) at 120 keV. The same area as the FIB area was recorded using the tiling option of the MAPS 3 software with the Ceta camera at 8500x magnification, resulting in a pixel size of 12Å. TEM imaging for melanosomes and transitioning leucosomes was done on an FEI Technai F20 equipped with a TEITZ XF416 detector, and recorded with the software SerailEM.

### Metabolomic Analysis

Dorsal fins were dissected from wild-type fish and *kita* mutants, ground into a powder with disposable pellet pestles under liquid nitrogen in pre-weighed tubes. Metabolites were extracted by adding 40 µL of ice cold methanol:acetonitrile:water (2:2:1) solution per milligram of tissue, followed by two freeze-thaw cycles that consisted of immersion in liquid nitrogen for 1 minute followed by incubation at 25°C for 10 seconds, sonication for 5 minutes at 25°C, and vortexing for 30 seconds. Samples were then incubated at −20°C for 1 h to precipitate protein and centrifuged at 20,000 x g for 10 minutes at 4°C. Following centrifugation, the supernatants were transferred to LC-MS vials for same-day analysis.

LC-MS based metabolomics was performed with a Vanquish Horizon UHPLC system (Thermo Scientific) coupled to a Q Exactive Plus Orbitrap mass spectrometer (Thermo Scientific). Metabolites were separated by using hydrophilic interaction liquid chromatography (HILIC) with an iHILIC-(P) Classic column (Hilicon) with same series guard column. Chromatographic solvents composed of A: water:acetonitrile (95:5), with the addition of 20 mM ammonium bicarbonate, 0.1% ammonium hydroxide, and 5 μM medronic acid; and B: acetonitrile:water (95:5) were utilized with the following gradient: 0 min, 90% B, 0.25 ml/min; 1 min, 90% B, 0.25 ml/min; 14 min, 25% B, 0.25 ml/min; 14.5 min, 25% B, 0.25 ml/min; 15 min, 90% B, 0.25 ml/min; 16.5 min, 90% B, 0.40 ml/min; 20 min, 90% B, 0.40 ml/min; 20.5 min, 90% B, 0.25 ml/min; 22 min, 90% B, 0.25 ml/min. The column was maintained at 40°C during analyses. For MS data acquisition, the ESI source was set at 3 kV for positive mode or 2.5 kV for negative mode; sheath gas 45; auxiliary gas 10; sweep gas 2; ion transfer tube 250°C; aux gas heater temperature 350°C.. The mass analyzer was set at a mass range 67 to 1000 Da; resolution was 120k for MS1 and 60k for MS/MS; MS/MSnormalized collision energy was set at 30 eV; maximum injection time 100 ms; and an isolation window of 1.5 Da. Raw data files were processed using Thermo Compound Discoverer 3.3 (Thermo Scientific), with an intensity threshold of 1E5, mass tolerance 5 ppm, and retention time tolerance of 0.2 min. Compounds were annotated by database searching and MS2 spectrum library matching using a cut-off of > 50.

### *In situ* Hybridization

Probe templates were amplified using Primestar-GXL (Takara) from cDNA prepared with SuperScript III (ThermoFisher) with the following primers: asip1 5’- AGAAGAGCAAGAAATCAGAAAAGAAG, 5’- aaaaTAATACGACTCACTATAGGTAATAATCATAAGGTGATTTTAAGTACCC; asip2b 5’- ATGACGACGGCGGTGCTGAAAGG, 5’- aaaaTAATACGACTCACTATAGGGTCTTCTTGGGGCAAAGGTGTCCC; bnc2 5’- ACTTAGTATTGAACAATCAGGTCTTAGAAT, 5’- aaaaTAATACGACTCACTATAGAGAAAGAAGAGTGTGTTATAGTTTTTGACT; fmodb 5’- attaaatatgtatacttgcaaaataaccag, 5’-aaaaTAATACGACTCACTATAGtctatagcatttgcttgaagatatagag; cdh11 5’-agatacaaccattgtgaaaatatcagtag, 5’- aaaaTAATACGACTCACTATAGatctttgactttgatattgactcgtacttt. Probes and tissue were prepared as described previously (46) with hybridization and post-hybridization washes performed on a BioLane HTI 16Vx platform (Intavis Bioanalytical Instruments) and post-staining sectioning in some cases as described (47).

### Single-cell RNA Sequencing and Analysis

We performed 3 single-cell RNA sequencing analyses, with each analysis using cells from wild-type *tyrp1b:PALM-mCherry* fish. All cells were sorted for cell membrane integrity by Dapi exclusion on a Sony SH800 fluorescence activated cell sorter. The first analysis, of melanophore–ML transcriptomes during normal development (dataset1_stages), used dorsal fins from fish of various sizes ranging from 9 mm standard length (SL) to 14 mm SL with sorting and retention of mCherry+ cells. The second analysis, to test differences in gene expression between fin regions (dataset2_positions), used dorsal fins from ∼10 mm SL fish that had been dissected into edge or non-edge regions (Fig. 2*D*), with retention of all cells to represent both mCherry+ and mCherry– populations. The third analysis, for acute effects of BMP inhibition, used cells from ∼12 mm SL fish treated with DMSO (vehicle control) or 2 µM LDN-193198 with DMSO for 48 hours (dataset3_BMPi), and also retained both mCherry+ and mCherry– populations after sorting. Cell dissociation, isolation and library production were performed as described (4) using a Chromium controller (10X Genomics) and single-cell 3’ solution V3 kit (10X Genomics). Quality control and quantification assays were performed using a Qubit fluorometer (ThermoFisher) and a 2100 Bioanalyzer (Agilent). Libraries were sequenced on an Illumina NextSeq 550 using 75-cycle, high output kits (read 1: 26 cycles, i7 Index: 8 cycles, read 2: 57 cycles).

We built a zebrafish STAR genome index using Lawson Lab zebrafish transcriptome annotation (48) plus manually annotated entries for mCherry transcript, filtered for protein-coding genes. Final cellular barcodes and Unique Molecular Identifiers (UMIs) were determined using Cell Ranger 6.0.1 (10X Genomics). Data were analyzed in Monocle3 (13, 14, 49) using methods described previously (1, 4). In brief, we filtered cells for quality to have less than 5% mitochondrial reads, and greater than 500 unique molecular identifiers and 200 genes expressed. Total numbers retained after filtering were 4,984 cells, 3,757 cells, and 14,866 cells for datasets1–3, respectively. Raw data and datasets in Monocle3 cell data set (cds) format are accessible by NCBI GEO (GSE282061). We used Uniform Manifold Approximation and Projection (UMAP) to project transcriptomic space in two dimensions followed by Louvain clustering, We assigned clusters to cell types by comparing genes detected to published cell-type specific markers (1, 4, 47) and transgenic lines (main text). Cell type marker analysis, differential expression and trajectory analyses used standard methods and functions as described (<u>cole-trapnell-lab.github.io/monocle3/docs/</u>). UMAP representation of melanophores–ML with trajectory inference used reduce_dimension() with umap.min.distance=0.35 and umap.n.neighbors=55, and cluster_cells() with resolution=1e-3. Gene module identification was performed using Monocle3 find_gene_modules() across clusters, and also as a function of pseudotime, with subsequent pathway analyses using Web-based Gene Set Analysis Toolkit (WEB-GESTALT; webgestalt.org) Identification of transcriptomic characteristics in cells expressing Agouti genes used *asip1* or *asip2b* expression values as parameters in Monocle3 fit_models(). Pathway signature scores were calculated as described (1) using marker loci compiled therein or curated manually from the literature.

For comparisons of pigment cell transcriptomes across types and anatomical locations [fin (this study) vs. body (1)] percentages of differentially expressed genes were calculated after down-sampling cell numbers to confer similar statistical powers across comparisons or cells were plotted in principal components space after generating a pseudo-bulk RNA-seq dataset comprising both fin and body data for analysis in DEseq2 (50).

## References

1. C. F. Kratochwil, R. Mallarino, Mechanisms Underlying the Formation and Evolution of Vertebrate Color Patterns. Annu Rev Genet 57, 135–156 (2023).

2. A. Brombin, E. E. Patton, Melanocyte lineage dynamics in development, growth and disease. Development 151 (2024).

3. D. M. Parichy, Evolution of pigment cells and patterns: recent insights from teleost fishes. Curr Opin Genet Dev 69, 88–96 (2021).

4. L. B. Patterson, D. M. Parichy, Zebrafish Pigment Pattern Formation: Insights into the Development and Evolution of Adult Form. Annu Rev Genet 53, 505–530 (2019).

5. S. Kondo, T. Miura, Reaction-diffusion model as a framework for understanding biological pattern formation. Science 329, 1616–1620 (2010).

6. J. P. Owen, R. N. Kelsh, C. A. Yates, A quantitative modelling approach to zebrafish pigment pattern formation. eLife 9 (2020).

7. A. Nakamasu, G. Takahashi, A. Kanbe, S. Kondo, Interactions between zebrafish pigment cells responsible for the generation of Turing patterns. Proc Natl Acad Sci U S A 106, 8429–8434 (2009).

8. D. M. Parichy, J. M. Turner, Temporal and cellular requirements for Fms signaling during zebrafish adult pigment pattern development. Development 130, 817–833 (2003).

9. V. M. Lewis et al., Fate plasticity and reprogramming in genetically distinct populations of Danio leucophores. Proc Natl Acad Sci U S A 116, 11806–11811 (2019).

10. J. Borovansky, M. Elleder, Melanosome degradation: fact or fiction. Pigment Cell Res 16, 280–286 (2003).

11. J. F. Rawls, S. L. Johnson, Zebrafish kit mutation reveals primary and secondary regulation of melanocyte development during fin stripe regeneration. Development 127, 3715–3724 (2000).

12. F. Figon et al., Catabolism of lysosome-related organelles in color-changing spiders supports intracellular turnover of pigments. Proc Natl Acad Sci U S A 118 (2021).

13. R. Tadokoro, Y. Takahashi, Intercellular transfer of organelles during body pigmentation. Curr Opin Genet Dev 45, 132–138 (2017).

14. R. Deis et al., Genetic control over biogenic crystal morphogenesis in zebrafish. Nat Chem Biol 10.1038/s41589-024-01722-1 (2024).

15. D. Gur et al., In situ differentiation of iridophore crystallotypes underlies zebrafish stripe patterning. Nat Commun 11, 6391 (2020).

16. N. W. Bellono, I. E. Escobar, A. J. Lefkovith, M. S. Marks, E. Oancea, An intracellular anion channel critical for pigmentation. eLife 3, e04543 (2014).

17. P. Wiriyasermkul, S. Moriyama, S. Nagamori, Membrane transport proteins in melanosomes: Regulation of ions for pigmentation. Biochim Biophys Acta Biomembr 1862, 183318 (2020).

18. H. B. Schonthaler et al., A mutation in the silver gene leads to defects in melanosome biogenesis and alterations in the visual system in the zebrafish mutant fading vision. Dev Biol 284, 421–436 (2005).

19. S. M. Mitchell, M. Graham, X. Liu, R. M. Leonhardt, Identification of critical amino acid residues in the regulatory N-terminal domain of PMEL. Scientific reports 11, 7730 (2021).

20. L. M. Saunders et al., Thyroid hormone regulates distinct paths to maturation in pigment cell lineages. eLife 8, e45181 (2019).

21. K. Petratou et al., A systems biology approach uncovers the core gene regulatory network governing iridophore fate choice from the neural crest. PLoS Genet 14, e1007402 (2018).

22. S. S. Lopes et al., Leukocyte tyrosine kinase functions in pigment cell development. PLoS Genet 4, e1000026 (2008).

23. M. M. Ollmann, M. L. Lamoreux, B. D. Wilson, G. S. Barsh, Interaction of Agouti protein with the melanocortin 1 receptor in vitro and in vivo. Genes Dev 12, 316–330 (1998).

24. M. Manceau, V. S. Domingues, R. Mallarino, H. E. Hoekstra, The developmental role of Agouti in color pattern evolution. Science 331, 1062–1065 (2011).

25. L. Cal et al., Countershading in zebrafish results from an Asip1 controlled dorsoventral gradient of pigment cell differentiation. Scientific reports 9, 3449 (2019).

26. L. Cal et al., Loss-of-function mutations in the melanocortin 1 receptor cause disruption of dorso-ventral countershading in teleost fish. Pigment Cell Melanoma Res 32, 817–828 (2019).

27. A. J. Aman et al., Transcriptomic profiling of tissue environments critical for post-embryonic patterning and morphogenesis of zebrafish skin. eLife 12 (2023).

28. Y. Liang, M. Grauvogl, A. Meyer, C. F. Kratochwil, Functional conservation and divergence of color-pattern-related agouti family genes in teleost fishes. J Exp Zool B Mol Dev Evol 336, 443–450 (2021).

29. L. Cal, P. Suarez-Bregua, J. M. Cerda-Reverter, I. Braasch, J. Rotllant, Fish pigmentation and the melanocortin system. Comp Biochem Physiol A Mol Integr Physiol 211, 26–33 (2017).

30. I. Braasch, J. H. Postlethwait, The teleost agouti-related protein 2 gene is an ohnolog gone missing from the tetrapod genome. Proc Natl Acad Sci U S A 108, E47–48 (2011).

31. C. Zhang et al., Pineal-specific agouti protein regulates teleost background adaptation. Proc Natl Acad Sci U S A 107, 20164–20171 (2010).

32. D. Wehner et al., Wnt/beta-catenin signaling defines organizing centers that orchestrate growth and differentiation of the regenerating zebrafish caudal fin. Cell reports 6, 467–481 (2014).

33. S. Stewart, A. W. Gomez, B. E. Armstrong, A. Henner, K. Stankunas, Sequential and opposing activities of Wnt and BMP coordinate zebrafish bone regeneration. Cell reports 6, 482–498 (2014).

34. R. Mateus et al., BMP Signaling Gradient Scaling in the Zebrafish Pectoral Fin. Cell reports 30, 4292–4302 e4297 (2020).

35. N. R. Infarinato et al., BMP signaling: at the gate between activated melanocyte stem cells and differentiation. Genes Dev 34, 1713–1734 (2020).

36. A. K. Gramann, A. M. Venkatesan, M. Guerin, C. J. Ceol, Regulation of zebrafish melanocyte development by ligand-dependent BMP signaling. eLife 8 (2019).

37. Q. Sun et al., Dedifferentiation maintains melanocyte stem cells in a dynamic niche. Nature 616, 774–782 (2023).

38. L. Vibert et al., An ongoing role for Wnt signaling in differentiating melanocytes in vivo. Pigment Cell Melanoma Res 30, 219–232 (2017).

39. Z. Zhang, K. Wu, Z. Ren, W. Ge, Genetic evidence for Amh modulation of gonadotropin actions to control gonadal homeostasis and gametogenesis in zebrafish and its noncanonical signaling through Bmpr2a receptor. Development 147 (2020).

40. J. Zhang et al., Repulsive guidance molecules b (RGMb): molecular mechanism, function and role in diseases. Expert Rev Mol Med 26, e24 (2024).

41. C. Siebold, T. Yamashita, P. P. Monnier, B. K. Mueller, R. J. Pasterkamp, RGMs: Structural Insights, Molecular Regulation, and Downstream Signaling. Trends Cell Biol 27, 365–378 (2017).

42. C. H. Bell et al., Structure of the repulsive guidance molecule (RGM)-neogenin signaling hub. Science 341, 77–80 (2013).

43. E. G. Healey et al., Repulsive guidance molecule is a structural bridge between neogenin and bone morphogenetic protein. Nat Struct Mol Biol 22, 458–465 (2015).

44. C. Simion, M. E. Cedano-Prieto, C. Sweeney, The LRIG family: enigmatic regulators of growth factor receptor signaling. Endocr Relat Cancer 21, R431–443 (2014).

45. S. van Erp et al., Lrig2 Negatively Regulates Ectodomain Shedding of Axon Guidance Receptors by ADAM Proteases. Dev Cell 35, 537–552 (2015).

46. H. M. Berns et al., Single-cell profiling of MC1R-inhibited melanocytes. Pigment Cell Melanoma Res 37, 291–308 (2024).

47. M. Lukoseviciute et al., From Pioneer to Repressor: Bimodal foxd3 Activity Dynamically Remodels Neural Crest Regulatory Landscape In Vivo. Dev Cell 47, 608–628 e606 (2018).

48. A. J. Thomas, C. A. Erickson, FOXD3 regulates the lineage switch between neural crest-derived glial cells and pigment cells by repressing MITF through a non-canonical mechanism. Development 136, 1849–1858 (2009).

49. K. Curran et al., Interplay between Foxd3 and Mitf regulates cell fate plasticity in the zebrafish neural crest. Dev Biol 344, 107–118 (2010).

50. N. Haupaix et al., The periodic coloration in birds forms through a prepattern of somite origin. Science 361 (2018).

51. C. F. Kratochwil et al., Agouti-related peptide 2 facilitates convergent evolution of stripe patterns across cichlid fish radiations. Science 362, 457–460 (2018).

52. A. A. Sharov et al., Bone morphogenetic protein (BMP) signaling controls hair pigmentation by means of cross-talk with the melanocortin receptor-1 pathway. Proc Natl Acad Sci U S A 102, 93–98 (2005).

53. M. Yaar et al., Bone morphogenetic protein-4, a novel modulator of melanogenesis. J Biol Chem 281, 25307–25314 (2006).

54. S. Y. A. Chow et al., Human sensory neurons modulate melanocytes through secretion of RGMB. Cell reports 40, 111366 (2022).

55. J. W. Lee et al., RUNX3 regulates cell cycle-dependent chromatin dynamics by functioning as a pioneer factor of the restriction-point. Nat Commun 10, 1897 (2019).

56. E. Nitzan, E. R. Pfaltzgraff, P. A. Labosky, C. Kalcheim, Neural crest and Schwann cell progenitor-derived melanocytes are two spatially segregated populations similarly regulated by Foxd3. Proc Natl Acad Sci U S A 110, 12709–12714 (2013).

57. Q. Shen, L. B. Toulabi, H. Shi, E. E. Nicklow, J. Liu, The forkhead transcription factor UNC-130/FOXD integrates both BMP and Notch signaling to regulate dorsoventral patterning of the C. elegans postembryonic mesoderm. Dev Biol 433, 75–83 (2018).

58. E. Nitzan et al., Dynamics of BMP and Hes1/Hairy1 signaling in the dorsal neural tube underlies the transition from neural crest to definitive roof plate. BMC Biol 14, 23 (2016).

59. R. Mevel, J. E. Draper, A. L. M. Lie, V. Kouskoff, G. Lacaud, RUNX transcription factors: orchestrators of development. Development 146 (2019).

60. Y. Ito, S. C. Bae, L. S. Chuang, The RUNX family: developmental regulators in cancer. Nat Rev Cancer 15, 81–95 (2015).

61. X. Zhang, L. Wang, X. Zeng, T. Fujita, W. Liu, Runx3 inhibits melanoma cell migration through regulation of cell shape change. Cell Biol Int 41, 1048–1055 (2017).

62. Y. Ito, K. Miyazono, RUNX transcription factors as key targets of TGF-beta superfamily signaling. Curr Opin Genet Dev 13, 43–47 (2003).

63. K. Petratou, S. A. Spencer, R. N. Kelsh, J. A. Lister, The MITF paralog tfec is required in neural crest development for fate specification of the iridophore lineage from a multipotent pigment cell progenitor. PLoS One 16, e0244794 (2021).

64. D. Arendt et al., The origin and evolution of cell types. Nat Rev Genet 17, 744–757 (2016).

65. I. K. Quigley et al., Pigment pattern evolution by differential deployment of neural crest and post-embryonic melanophore lineages in Danio fishes. Development 131, 6053–6069 (2004).

66. D. M. Parichy, M. R. Elizondo, M. G. Mills, T. N. Gordon, R. E. Engeszer, Normal table of postembryonic zebrafish development: staging by externally visible anatomy of the living fish. Developmental Dynamics 238, 2975–3015 (2009).

## SI References

1 L. M. Saunders et al., Thyroid hormone regulates distinct paths to maturation in pigment cell lineages. eLife 8, e45181 (2019).

2 Y. Usui, S. Kondo, M. Watanabe, Melanophore multinucleation pathways in zebrafish. Dev Growth Differ 60, 454–459 (2018).

3 F. Figon et al., Catabolism of lysosome-related organelles in color-changing spiders supports intracellular turnover of pigments. Proc Natl Acad Sci U S A 118 (2021).

4 V. M. Lewis et al., Fate plasticity and reprogramming in genetically distinct populations of Danio leucophores. Proc Natl Acad Sci U S A 116, 11806–11811 (2019).

5 M. Goda, A. Miyagi, T. Kitamoto, M. Kondo, H. Hashimoto, Uric acid is a major chemical constituent for the whitish coloration in the medaka leucophores. Pigment Cell Melanoma Res 36, 416–422 (2023).

6 B. M. McCluskey, S. Uji, J. L. Mancusi, J. H. Postlethwait, D. M. Parichy, A complex genetic architecture in zebrafish relatives Danio quagga and D. kyathit underlies development of stripes and spots. PLoS Genet 17, e1009364 (2021).

7 Q. Sun, Q. Hao, K. V. Prasanth, Nuclear Long Noncoding RNAs: Key Regulators of Gene Expression. Trends Genet 34, 142–157 (2018).

8 S. Das, C. Pradhan, D. Pillai, beta-Defensin: An adroit saviour in teleosts. Fish Shellfish Immunol 123, 417–430 (2022).

9 X. C. Yuan, Y. X. Tao, Ligands for Melanocortin Receptors: Beyond Melanocyte-Stimulating Hormones and Adrenocorticotropin. Biomolecules 12 (2022).

10 S. I. Candille et al., A -defensin mutation causes black coat color in domestic dogs. Science 318, 1418–1423 (2007).

11 K. D. Poss, Advances in understanding tissue regenerative capacity and mechanisms in animals. Nat Rev Genet 11, 710–722 (2010).

12 J. Diener, L. Sommer, Reemergence of neural crest stem cell-like states in melanoma during disease progression and treatment. Stem Cells Transl Med 10, 522–533 (2021).

13 J. Cao et al., The single-cell transcriptional landscape of mammalian organogenesis. Nature 566, 496–502 (2019).

14 X. Qiu et al., Reversed graph embedding resolves complex single-cell trajectories. Nat Methods 14, 979–982 (2017).

15 K. W. Lee et al., RCHY1 and OPTN are required for melanophagy, selective autophagy of melanosomes. Proc Natl Acad Sci U S A 121, e2318039121 (2024).

16 K. Petratou, S. A. Spencer, R. N. Kelsh, J. A. Lister, The MITF paralog tfec is required in neural crest development for fate specification of the iridophore lineage from a multipotent pigment cell progenitor. PLoS One 16, e0244794 (2021).

17 R. Deis et al., Genetic control over biogenic crystal morphogenesis in zebrafish. Nat Chem Biol 10.1038/s41589-024-01722-1 (2024).

18 C. W. Higdon, R. D. Mitra, S. L. Johnson, Gene expression analysis of zebrafish melanocytes, iridophores, and retinal pigmented epithelium reveals indicators of biological function and developmental origin. PLoS ONE 8, e67801 (2013).

19 H. S. Jang et al., Epigenetic dynamics shaping melanophore and iridophore cell fate in zebrafish. Genome Biol 22, 282 (2021).

20 E. S. Mo, Q. Cheng, A. V. Reshetnyak, J. Schlessinger, S. Nicoli, Alk and Ltk ligands are essential for iridophore development in zebrafish mediated by the receptor tyrosine kinase Ltk. Proc Natl Acad Sci U S A 114, 12027–12032 (2017).

21 S. S. Lopes et al., Leukocyte tyrosine kinase functions in pigment cell development. PLoS Genet 4, e1000026 (2008).

22 D. M. Parichy et al., Mutational analysis of endothelin receptor b1 (rose) during neural crest and pigment pattern development in the zebrafish Danio rerio. Dev Biol 227, 294–306 (2000).

23 P. Salis et al., Developmental and comparative transcriptomic identification of iridophore contribution to white barring in clownfish. Pigment Cell Melanoma Res 32, 391–402 (2019).

24 E. Moro et al., In vivo Wnt signaling tracing through a transgenic biosensor fish reveals novel activity domains. Dev Biol 366, 327–340 (2012).

25 C. L. Stoick-Cooper et al., Distinct Wnt signaling pathways have opposing roles in appendage regeneration. Development 134, 479–489 (2007).

26 A. J. Aman, A. N. Fulbright, D. M. Parichy, Wnt/beta-catenin regulates an ancient signaling network during zebrafish scale development. eLife 7, 10.7554/eLife.37001 (2018).

27 A. K. Gramann, A. M. Venkatesan, M. Guerin, C. J. Ceol, Regulation of zebrafish melanocyte development by ligand-dependent BMP signaling. eLife 8 (2019).

28 B. M. McCluskey, J. H. Postlethwait, Phylogeny of zebrafish, a “model species,” within Danio, a “model genus”. Mol Biol Evol 32, 635–652 (2015).

29 D. M. Parichy, Advancing biology through a deeper understanding of zebrafish ecology and evolution. eLife 4, e05635 (2015).

30 S. He et al., Molecular phylogenetics of the family Cyprinidae (Actinopterygii: Cypriniformes) as evidenced by sequence variation in the first intron of S7 ribosomal protein-coding gene: further evidence from a nuclear gene of the systematic chaos in the family. Mol Phylogenet Evol 46, 818–829 (2008).

31 R. L. Mayden et al., Phylogenetic relationships of Danio within the order Cypriniformes: a framework for comparative and evolutionary studies of a model species. J Exp Zoolog B Mol Dev Evol 308, 642–654 (2007).

32 R. Mevel, J. E. Draper, A. L. M. Lie, V. Kouskoff, G. Lacaud, RUNX transcription factors: orchestrators of development. Development 146 (2019).

33 Y. Ito, S. C. Bae, L. S. Chuang, The RUNX family: developmental regulators in cancer. Nat Rev Cancer 15, 81–95 (2015).

34 D. M. Parichy, J. F. Rawls, S. J. Pratt, T. T. Whitfield, S. L. Johnson, Zebrafish sparse corresponds to an orthologue of c-kit and is required for the morphogenesis of a subpopulation of melanocytes, but is not essential for hematopoiesis or primordial germ cell development. Development 126, 3425–3436 (1999).

35 J. E. Spiewak et al., Evolution of Endothelin signaling and diversification of adult pigment pattern in Danio fishes. PLoS Genet 14, e1007538 (2018).

36 S. K. McMenamin et al., Thyroid hormone-dependent adult pigment cell lineage and pattern in zebrafish. Science 345, 1358–1361 (2014).

37 R. E. Engeszer, L. B. Patterson, A. A. Rao, D. M. Parichy, Zebrafish in the wild: a review of natural history and new notes from the field. Zebrafish 4, 21–40 (2007).

38 K. Hoshijima et al., Highly Efficient CRISPR-Cas9-Based Methods for Generating Deletion Mutations and F0 Embryos that Lack Gene Function in Zebrafish. Dev Cell 51, 645–657 e644 (2019).

39 D. M. Parichy, M. R. Elizondo, M. G. Mills, T. N. Gordon, R. E. Engeszer, Normal table of postembryonic zebrafish development: staging by externally visible anatomy of the living fish. Developmental Dynamics 238, 2975–3015 (2009).

40 K. M. Kwan et al., The Tol2kit: A multisite gateway-based construction kit forTol2 transposon transgenesis constructs. Developmental Dynamics 236, 3088–3099 (2007).

41 E. Provost, J. Rhee, S. D. Leach, Viral 2A peptides allow expression of multiple proteins from a single ORF in transgenic zebrafish embryos. Genesis 45, 625–629 (2007).

42 M. L. Suster, H. Kikuta, A. Urasaki, K. Asakawa, K. Kawakami, Transgenesis in zebrafish with the tol2 transposon system. Methods Mol Biol 561, 41–63 (2009).

43 C. Kizilyaprak, Y. D. Stierhof, B. M. Humbel, Volume microscopy in biology: FIB-SEM tomography. Tissue Cell 57, 123–128 (2019).

44 C. Kizilyaprak, G. Longo, J. Daraspe, B. M. Humbel, Investigation of resins suitable for the preparation of biological sample for 3-D electron microscopy. J Struct Biol 189, 135–146 (2015).

45 J. R. Kremer, D. N. Mastronarde, J. R. McIntosh, Computer visualization of three-dimensional image data using IMOD. J Struct Biol 116, 71–76 (1996).

46 I. K. Quigley et al., Pigment pattern evolution by differential deployment of neural crest and post-embryonic melanophore lineages in Danio fishes. Development 131, 6053–6069 (2004).

47 A. J. Aman et al., Transcriptomic profiling of tissue environments critical for post-embryonic patterning and morphogenesis of zebrafish skin. eLife 12 (2023).

48 N. D. Lawson et al., An improved zebrafish transcriptome annotation for sensitive and comprehensive detection of cell type-specific genes. eLife 9 (2020).

49 C. Trapnell et al., The dynamics and regulators of cell fate decisions are revealed by pseudotemporal ordering of single cells. Nat Biotechnol 32, 381–386 (2014).

50 M. I. Love, W. Huber, S. Anders, Moderated estimation of fold change and dispersion for RNA-seq data with DESeq2. Genome Biol 15, 550 (2014).

